# Mitochondrial genome analyses uncover intriguing population structure across the invasive trajectory of an iconic invader

**DOI:** 10.1101/2023.09.17.558141

**Authors:** K Cheung, TG Amos, R Shine, JL DeVore, S Ducatez, RJ Edwards, LA Rollins

## Abstract

Invasive species offer insights into rapid adaptations in novel environments. The iconic cane toad (*Rhinella marina*) is an excellent model for studying rapid adaptation during invasion. Previous research using the mitochondrial NADH dehydrogenase 3 (*ND3*) gene in the Hawai’ian and Australian invasive populations found a single haplotype, indicating an extreme genetic bottleneck following introduction. Nuclear genetic diversity also exhibited reductions across the genome in these two populations. Here, we investigated the mitochondrial genome diversity of cane toads across this invasion trajectory. We created the first reference mitochondrial genome of the cane toad with long-read sequencing and constructed a phylogeny of Anura full mitochondrial genomes. We used transcriptomic data of 125 individuals from the native (French Guiana) and introduced (Hawai’i and Australia) ranges to construct nearly-complete mitochondrial genomes for population genomics analyses. As expected, the cane toad belongs to family Bufonidae, distinct from genus *Bufo*. In agreement with previous investigations of these populations, we identified genetic bottlenecks in both Hawai’ian and Australian introduced populations, alongside evidence of population expansion in the invasive ranges. Although mitochondrial genetic diversity in introduced populations was reduced, our results revealed that it had been underestimated: we identified 45 mitochondrial haplotypes in Hawai’ian and Australian samples, none of which were found in the native range. Additionally, we identified two distinct groups of haplotypes from the native range, separated by a minimum of 110 base pairs (0.6%). These findings enhance our understanding of Anura phylogenetics and how invasion has shaped the genetic landscape of this species.

## Introduction

Invasive species are a substantial global concern, particularly due to likely range expansions under climate change (Bonebrake et al., 2018; Seebens et al., 2017), an increasing rate of anthropogenic introductions (Lowe, Browne, Boudjelas, & De Poorter, 2000), and their negative impacts on native biodiversity and the economy (Bradshaw et al., 2016; Mainka & Howard, 2010). Invasive species often rapidly adapt to their new environments (Rollins et al., 2013), creating an opportunity to study the process of rapid evolution. Biological invasions sometimes present a genetic paradox where, despite small founding population sizes that often reduce genetic diversity in the introduced range (i.e. genetic bottlenecks), they still maintain the ability to readily adapt to novel conditions (Allendorf & Lundquist, 2003; Estoup et al., 2016; Schrieber & Lachmuth, 2017).

Various genetic markers, including those from the nuclear and mitochondrial genomes, have been employed to investigate the introduction history and evolutionary processes of invasive species. Because of their distinct mode of inheritance and mutation rates, nuclear and mitochondrial markers may yield different insights into the evolutionary events underlying invasion (Toews & Brelsford, 2012). Genetic variation in invasive species is shaped by complex interplays among genetic drift, demography and selection. Small founding populations, common during an invasion, can promote genetic drift, leading to reduced genetic diversity. Furthermore, whilst genetic drift results in less efficient selection (Gravel, 2016), populations expanding their range into new environments may experience novel selection regimes (Sakai et al., 2001) and spatial sorting (Shine, Brown, & Phillips, 2011) that further reduce genetic diversity.

Mitochondrial DNA (mtDNA) is more susceptible to genetic drift than nuclear markers due to its uniparental inheritance and small effective population size (Klucnika & Ma, 2019; Xiao, Nguyen, Wu, & Hao, 2017). Nonetheless, mtDNA has been extensively used to study the evolutionary history of different species or populations within the same species because of its non-recombining nature and purported near-neutrality. However, the neutrality of mtDNA has been challenged by evidence of selection, likely due to the important role of mitochondria in ATP production and cellular activities (Meiklejohn, Montooth, & Rand, 2007). Deleterious mutations are rapidly removed by purifying selection to maintain the function of the electron transport chain. Further, variation in mtDNA has been linked to adaptation to high latitudes (Chen et al., 2020; Wang et al., 2021) and temperature (Baker et al., 2019; Cheng, Liu, Yang, Sun, & Kong, 2013; Silva, Lima, Martel, & Castilho, 2014). There exists a substantial repository of mtDNA data in public databases, enabling comprehensive comparative genetic analyses, increasing its usefulness as a marker.

The cane toad (*Rhinella marina*) is one of the world’s most infamous, and well-studied, invasive species (Lowe et al., 2000). Cane toads are native to Central and South America (Zug & Zug, 1979) and have been introduced to multiple Caribbean islands for biological control of cane beetles, a parasite of sugarcane (Easteal, 1981). Cane toads were transported from the Caribbean to Hawai’i in 1932 for cane beetle control and, in 1935, 101 Hawai’ian cane toads were translocated from Oahu to Queensland, Australia (Turvey, 2013). However, they failed to control cane beetles in Australia and, facilitated by a high reproductive rate, subsequently spread across the continent to arid Western Australia. This species’ inexorable spread throughout its introduced range in Australia has had catastrophic effects on the naïve predators it has encountered (Chinchio et al., 2020). Freshwater crocodiles, snakes, lizards, and quolls have been particularly badly affected by the cane toad invasion in Australia (Shine, 2010). The colonization of the cane toad is still on-going; toads occupy over 1.2 million km^2^ of Australia (Urban, Phillips, Skelly, & Shine, 2008) including tropical Queensland, the Northern Territory, and the Kimberley region of northern Western Australia (Radford et al., 2019).

Like in many other invasions, the cane toads’ translocation resulted in genetic bottlenecks. Australian cane toads have very low MHC class I and class II (Lillie, Cui, Shine, & Belov, 2016; Lillie, Dubey, Shine, & Belov, 2017; Lillie, Shine, & Belov, 2014) and microsatellite (Leblois, Rousset, Tikel, Moritz, & Estoup, 2000) diversity. Previous analysis of the NADH dehydrogenase 3 (*ND3*) mitochondrial gene from 31 individuals sampled from Hawai’i and Australia identified a single haplotype, a substantial reduction of haplotype diversity compared to the native range (Slade & Moritz, 1998), further supporting the presence of genetic bottlenecks in the invasive ranges. Despite this low genetic variation, phenotypic traits differ substantially across the cane toad’s Australian range including immune function, dispersal ability, and behaviour (Alford, Brown, Schwarzkopf, Phillips, & Shine, 2009; G. P. Brown, Phillips, Dubey, & Shine, 2015; Gruber, Brown, Whiting, & Shine, 2018; Gruber, Whiting, Brown, & Shine, 2017; Phillips, Brown, & Shine, 2010). Some of these traits have been shown to be intergenerationally transmitted, consistent with genetic causes. For example, toads from range-core and range-edge populations that were raised in a common-garden experiment showed divergence of behavioural traits and morphology (Gruber, Brown, Whiting, & Shine, 2017; Hudson, Brown, & Shine, 2016). The invasion success of this species in Australia, coupled with evidence of evolutionary change since introduction make it an ideal system for the study of rapid evolution during invasion.

Despite extensive research into toad evolution in the introduced range, we need improved genetic resources to clarify the molecular mechanisms underlying these changes. Transcriptome data from spleen, muscle and/or brain tissues of toads from French Guiana, Hawai’i, and Australia, and liver tissue from Brazil exist (Richardson et al., 2018; Rollins, Richardson, & Shine, 2015; Selechnik, Richardson, Shine, Brown, & Rollins, 2019; Yagound et al., 2022) and a draft reference genome (Edwards et al., 2018) is available, but population genomic analyses of cane toads across this invasion trajectory have thus far been restricted to a single study of single nucleotide polymorphisms (SNPs) from the nuclear genome (Selechnik, Richardson, Shine, DeVore, et al., 2019). Here, we present a complete cane toad mitochondrial genome (mitogenome) and explore the species’ phylogenetic relationships with other Anurans. We then investigate evolution across the invasion trajectory using mtDNA data derived from RNASeq and whole-genome resequencing. We compare our data to those of a previously-published study of *ND3* on the same populations (Slade & Moritz, 1998), to determine whether our whole-mitochondrial genome analysis can extend population-genetic inferences. We predicted that there would be decreasing genetic diversity within the mitochondrial genome with increasing distance from the native range due to genetic bottlenecks. We also predicted that we would identify more genetic diversity within Australia than has been previously characterised, given the phenotypic evidence of adaptation to this novel environment.

## Materials and Methods

### Reference mitochondrial genome

High-molecular-weight genomic DNA was extracted from the liver of a female cane toad collected in the Kimberley region of Western Australia and sequenced on Illumina HiSeq X Ten and PacBio RS II platforms as described previously (Edwards et al., 2018). The mitogenome was assembled from 36 SMRT cells generated from the first two SMRTBell libraries (ENA experiments ERX2389178 and ERX2389179).

The 17,757 basepair (bp) *Bufo japonicus* complete mitochondrial genome (gi:157939553) was downloaded from NCBI on 29/3/2016 and used to identify mitochondrial subreads from the raw sequencing data. GABLAM (Davey, Shields, & Edwards, 2006) was used to map the *B. japonicus* mtDNA onto PacBio subreads using blastn (blast+ v2.2.31) and extract 143 subreads that had 75%+ bidirectional coverage, *e.g.* at least 75% of the subread mapped onto the mtDNA and at least 75% of the mtDNA was covered by the subread. Any subreads that mapped the negative strand were reverse complemented. The subread with the best global hit to the *B. japonicus* mtDNA was selected as the core of the cane toad mitochondrial genome. A second GABLAM search was performed of the 143 subreads against this new query, retaining 29 sequences that had >80% global sequence identity within local matches in the same orientation and order, excluding any “wrapping” due to circularity. These 29 sequences were aligned using MAFFT v2.273 (Katoh & Standley, 2013) with a very small gap penalty to account for the high indel rate (--localpair --maxiterate 1000 --op 0.1 --ep 0.1 --lop -0.1 --nuc). SeqSuite (Edwards, Paulsen, Aguilar Gomez, & Perez-Bercoff, 2020) was used to generate a consensus sequence from the alignment using the most frequent non-gap base call at each position. Any column with fewer than 5 non-gap sequences was excluded from the consensus. All 143 raw subreads were mapped onto the consensus using BLASR and the sequence was polished using Quiver (SMRT Analysis v2.3.0). GABLAM was used to identify the overlapping ends of the sequence, which was then circularised by trimming off the first 1957 bp for a final length of 18,152 bp. A final polishing step was performed using the Illumina data (ENA experiment ERX2845325), mapped with Bowtie v2.3.3 (Langmead & Salzberg, 2012) and error-corrected with Pilon v1.20 (Walker et al., 2014).

The mitochondrial genome was annotated using MITOS online (Bernt et al., 2013) with the vertebrate mtDNA genetic code (Genetic Code Table 02). Annotated protein-coding genes were manually curated and extended to include stop codons where required. Where no canonical in-frame stop codon was found, it was assumed that a partial (T or TA) stop codon would be completed upon polyadenylation. Codon usage was calculated using https://www.bioinformatics.org/sms2/codon_usage.html (Accessed: 2023-03-29). The final annotated mitochondrial genome was uploaded to NCBI (Accession number: NC_066225.1). The mitogenome map was drawn using the CGView server (http://stothard.afns.ualberta.ca/cgview_server/).

### Phylogenetic analysis

Complete mitochondrial genomes of 203 Anura, and 5 non-Anura outgroup species from Caeciliidae and Caudata), were downloaded from NCBI BioProject database (Table S3, Downloaded 2022-09-11). Each mitochondrial genome was re-circularised and edited to end with the control region for subsequent analyses. The multiple sequence alignment was conducted with the cane toad reference mitogenome using MAFFT with auto mode to find the best alignment algorithm (Katoh & Standley, 2013). The resulting alignment was visually inspected using AliView (Larsson, 2014) without further trimming. Maximum likelihood phylogenetic analysis was conducted using IQ-Tree multicore version 2.2.2.7 (default parameters, ultrafast bootstrap (UFBoot) and SH-aLRT branch test with 10000 replicates) (Minh et al., 2020). Clades are considered reliable if SH-aLRT value >= 80% and UFboot value >=95%. The best fit model was found by ModelFinder embedded in IQ-Tree (Kalyaanamoorthy, Minh, Wong, von Haeseler, & Jermiin, 2017). The phylogenetic tree was visualised and edited using R package *ggtree* (Yu, Smith, Zhu, Guan, & Lam, 2017).

### Sequence data for population- and species-level analyses

Previously published cane toad RNASeq data from different tissues were retrieved and downloaded from NCBI BioProject PRJNA510261 and PRJNA395127 (spleen), PRJNA479937 (brain) and PRJNA277985 (muscle) for analysis. A total of 125 individuals with 144 tissue samples were included, sampled in French Guiana (*n* = 24), Hawai’i (*n* = 8) and Australia (*n* = 93) (Figure 1; Table S1). For some individuals, RNASeq data from both spleen and brain were available; in such cases, we used data from brains. Raw reads were processed as described previously with STAR v2.7.2b (Dobin et al., 2013) and Genome Analysis Toolkit (GATK) v4.1.9.0 (McKenna et al., 2010), using the extracted mitogenome from the draft genome as a reference (Selechnik, Richardson, Shine, DeVore, et al., 2019).

**Figure 1:**
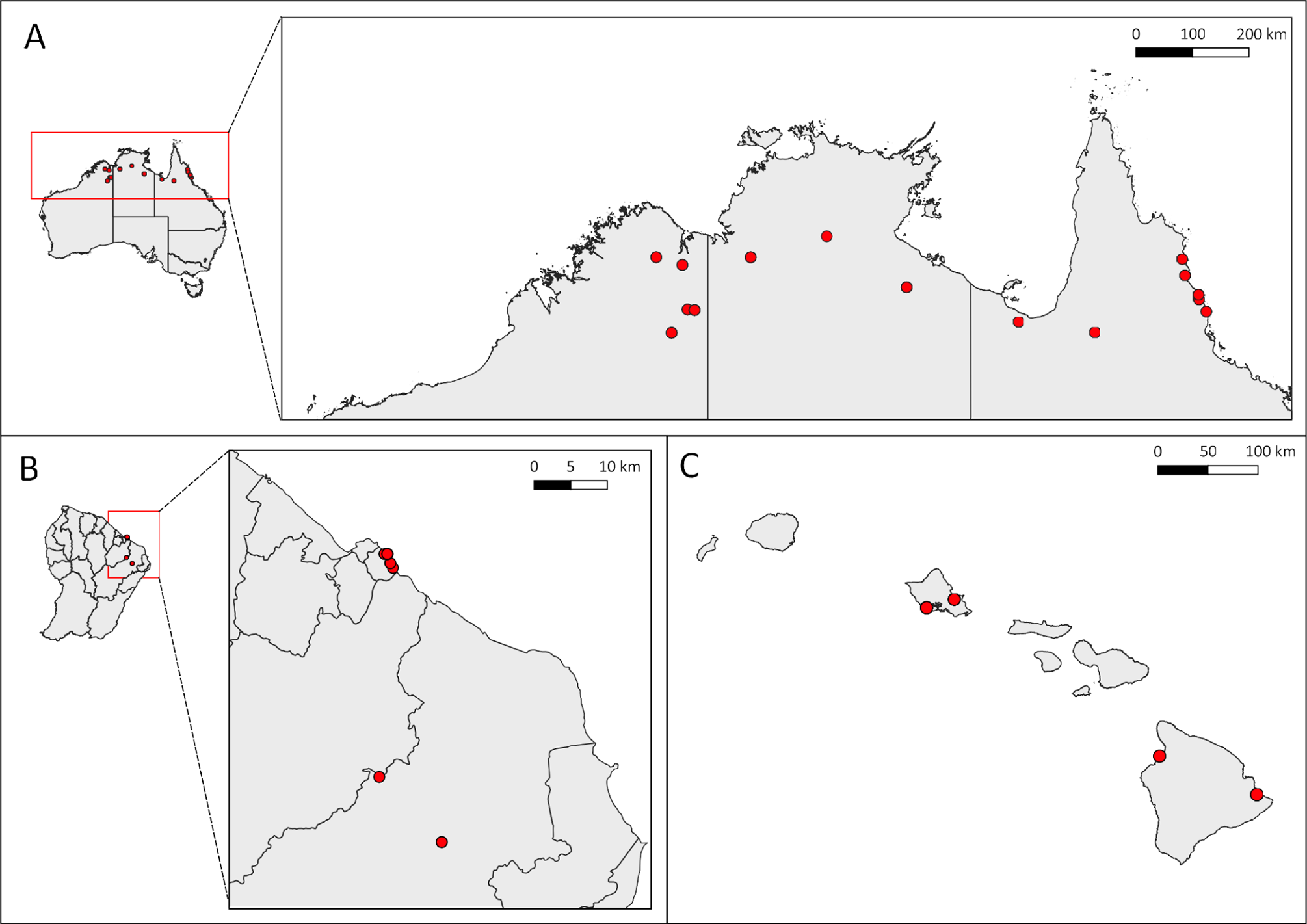
Sampling locations of the cane toad (*Rhinella marina*) in (A) Australia and (B) French Guiana (the native range) and (C) Hawai’i. Red dots in French Guiana, Hawai’i and Australia indicate sample collection sites.

Genomic DNA was extracted from fifteen cane toad samples, including two from toe clips, two from spleen, four from liver and seven from muscle tissues using a Gentra PureGene DNA extraction kit (QIAGEN) (Table S1).

For two samples (RMH006 and RM265), paired-end sequencing libraries were prepared using the TruSeq DNA PCR-free kit (Illumina). These libraries were sequenced on the Illumina NovaSeq 6000 platform at the Ramaciotti Centre for Genomics, University of New South Wales, Sydney, Australia (UNSW Sydney). The remaining thirteen samples had paired-end sequencing libraries constructed according to the manufacturer’s specifications for the NEBNext Ultra DNA Library Prep Kit for Illumina (NEB). Three samples (RMF031, RMF042 and RMF044) were sequenced on the 10X Genomics Chromium platform at the Ramaciotti Centre for Genomics, UNSW Sydney. The remaining libraries were sequenced on the Illumina NovaSeq 6000 platform at Deakin Genomic Centre, Victoria, Australia with a target depth of 20-fold coverage per genome. Raw sequencing reads from each sample underwent initial processing to remove adapter sequences. Trimmomatic v0.38 (Bolger, Lohse, & Usadel, 2014) was employed using the following parameters: ILLUMINACLIP:TruSeq3-PE.fa:2:30:10:4 SLIDINGWINDOW:5:30 AVGQUAL:30 MINLEN:36.

Mitochondrial segments, *ND3* gene with tRNA-Gly and tRNA-Arg flanking sequence (*ND3* dataset hereafter) were obtained from previous studies (Acevedo, Lampo, & Cipriani, 2016; Slade & Moritz, 1998) (Table S1). We aligned our data with this *ND3* dataset using the R package DECIPHER (Wright, 2016) and trimmed our data to match the existing sequences.

### Construction of mtDNA genome and variant calling

GATK HaplotypeCaller (Poplin et al., 2018) was used to call SNPs and indels using the following criteria: soft clipped bases were not included and variants with minimum Phred-scaled confidence of 20 threshold were kept. To filter low quality variants, GATK VariantFiltration (Van der Auwera, 2020) was used to remove variants in the resulting VCF file: clusters of 3 SNPs within a window of 35 bases, variants with Fisher Strand (FS) greater than 30.0, QualbyDepth (QD) less than 2.0, depth of coverage (DP) less than 20.0 and allelic frequency (AF) less than 0.05. Due to the difficulty of mapping short-read RNASeq data to sequences with multiple repeats, and the risk of introducing false-positive variant calls, we excluded variants from the repetitive sequences in the hypervariable region 2 (HV2) of the mitochondrial control region. We determined the consensus sequence of each sample using FastaAlternateReferenceMaker (McKenna et al., 2010).

Consensus mitogenome sequences from individuals with both RNASeq and WGS data were aligned against each other using DECIPHER R packages (Wright, 2016, 2020) using default settings. The alignments were further visualised and analysed with AliView (Larsson, 2014).

### NUMT identification

The presence of NUMTs in the cane toad’s genome has not been documented previously. The cane toad draft genome was retrieved and downloaded from NCBI BioProject PRJEB24695. We used RepeatModeler (http://www.repeatmasker.org/RepeatModeler/) to construct species-specific repeat library and masked the genome using RepeatMasker (Tarailo-Graovac & Chen, 2009). NUMTFinder v0.5.3 (Edwards et al., 2021) was used to identify potential NUMTs by searching the full and repeat-masked versions of the nuclear genome with our new mtDNA assembly.

### Population genetics analyses

The spatial distribution of haplotypes was examined by creating a median-joining haplotype network using Network and postprocessed using the maximum parsimony calculation to remove unnecessary median vectors and links (Polzin & Vahdati Daneshmand, 2003). The final network was drawn using Network Publisher v2.1.2.5 (Fluxus Engineering, Clare, UK). This same procedure also was used to analyse the *ND3* dataset.

We then investigated population genetic diversity and differentiation. The following diversity indices were calculated using DnaSP v6 (Rozas et al., 2017): nucleotide diversity (π), number of polymorphic sites, number of parsimony informative sites, number of haplotypes (*H*) and haplotype diversity (*h*). Due to the uneven sample size across populations, haplotype richness (*HR*) was calculated using FSTAT v2.9.4 (Goudet, 1995) to measure the number of alleles independent of the sample size. DnaSP was also used to generate a haplotype list for Arlequin software v3.5.2.2 (Excoffier & Lischer, 2010) and a Roehl data file for use with Network software 10.2.0.0 (Bandelt, Forster, & Rohl, 1999; Bandelt, Forster, Sykes, & Richards, 1995). To study the population structure, Arlequin was used to calculate genetic differentiation among and between populations by pairwise fixation indices (*Fst*) and Analysis of Molecular Variance (AMOVA) to estimate the genetic variation among and within populations.

To study the demographic history of each population, we used both neutrality statistics and mismatch distribution tests to infer population expansion. Fu’s *Fs* statistics was produced using Arlequin and Ramos-Onsins and Rozas *R_2_* statistics were produced using DnaSP. Fu’s *Fs* test calculates the probability of observing a number of haplotype similar or smaller than the observed number of samples (Fu, 1997). Both tests are based on the infinite-site model without recombination, which is suitable for analysing mitochondrial genomic data. A significant negative value of *Fs* indicates an excess of rare alleles and haplotypes due to a sudden population growth after a population bottleneck or genetic hitchhiking while positive values indicate a lack of rare mutations, resulting from balancing selection or population stability. Ramos-Onsins and Rozas *R_2_* statistic compares the difference between the number of singleton mutations and the average number of segregating sites (Ramos-Onsins & Rozas, 2002). The *R_2_* statistic is known to have better power than Fu’s *Fs* when the sample size is small (< 10) or the number of segregating sites is low (Ramirez-Soriano, Ramos-Onsins, Rozas, Calafell, & Navarro, 2008). A small positive value of *R_2_* indicates population expansion. The statistical power of these tests was validated using 10,000 simulations. Mismatch distributions of the frequency of pairwise differences between haplotype pairs in each population were also examined using the sudden expansion and spatial expansion models implemented in Arlequin (Excoffier, 2004; Schneider & Excoffier, 1999). The sum of square deviations (SSD) was used to test the differences between observed and expected values while the raggedness index was used to explore the smoothness of the observed mismatch distribution. The statistical significance of these tests was validated with 500 replicates in Arlequin. A non-significant SSD value and a small raggedness index value with a unimodal distribution indicates no deviation of the observed data from the expectation under the expansion model.

## Results

### Summary of the cane toad reference mitochondrial genome

The mitogenome of *R. marina* (Figure 2) consists of 18,152 bp, including 22 tRNA genes, 2 rRNA genes and 13 protein-coding genes, with 28 genes located on the forward (+) strand and 9 located on reverse (-) strand (Table S2). There is also a 2,753 bp GC-rich control region containing the origin of replication. Two groups of tandem repeats were observed in the latter part of the control region: 530 bps with 6-full repeats and 350 bps with 3-full repeats. The gene order of the cane toad mitogenome was examined by aligning with the members of its sister clades (*Bufo* and *Bufotes*) and we found no gene rearrangements. The base composition of the mitogenome was A: 29%, T: 33%, G: 14% and C: 24%, with a bias with 62% of A + T. The codon usages of those with nucleotide G in the third codon position were lower than those with A and T. GCG (Ala) was the least used codon and TGG was the only codon used for encoding Tryptophan. All protein-coding genes used ATG as start codon.

**Figure 2:**
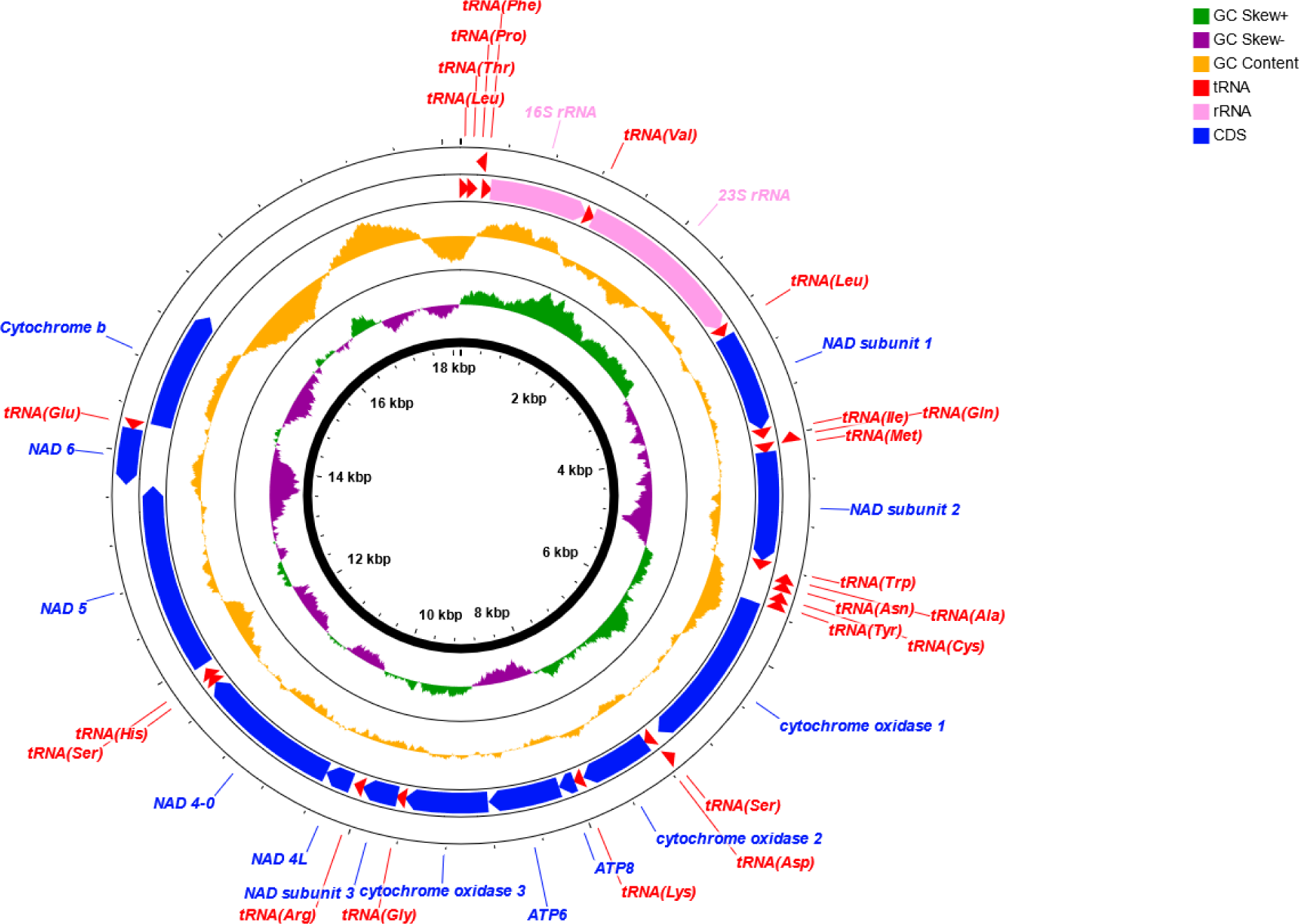
Mitochondrial genome structure of *R. marina*. Genes, GC skew and GC content were shown in the inner circle, middle circle and outer circle respectively. The innermost circle indicates the negative strand and the outer circle indicates the positive strand. Genes are coloured by type: red, tRNA; pink, rRNA; blue, coding sequence. Starting and stop positions of each gene are described in Table S3.

### Phylogenetics analysis

We produced a maximum likelihood phylogenetic tree using whole mitogenomes of Anura. The best fit model was GTR+F+R9 according to lowest Bayesian information Criterion (BIC). The location of *R. marina* in the phylogeny of Anura is shown in Figure 3. *Rhinella marina* was grouped with *Anaxyrus americanus*, an American Bufonid and previously classified under *Bufo* genus, with 100% statistical support in both UFBoot and SH-aLRT in branch tests. Four species of *Bufotes* and four of *Bufo* were well resolved into two subclades. Even though there was high branch support for differentiation between the genera *Rhinella* and *Anaxyrus*, and high support for species within *Bufotes* and *Bufo*, the divergence among genera was not significantly supported, which may indicate the signals in the whole mitochondrial genomes were insufficient for resolving these clades or that wider taxonomic coverage is required to resolve this phylogeny.

**Figure 3:**
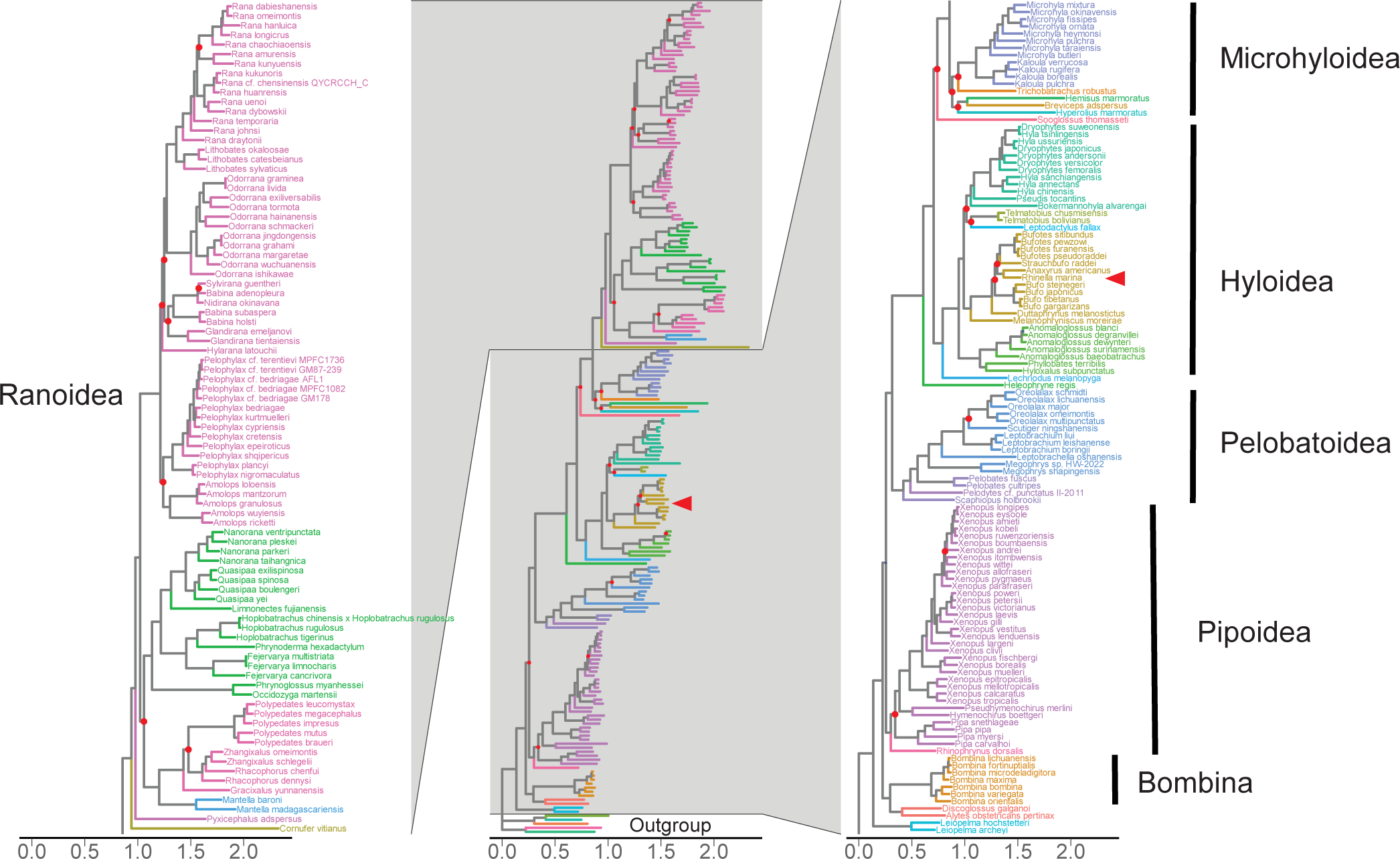
Maximum-likelihood phylogenetic tree of the complete mitochondrial genomes of 203 Anuran species downloaded from the NCBI BioProject public database. Five non-Anuran species from Caudata and Caeciliidae were used as outgroup (Table S3). The dendrogram was constructed using IQ-Tree multicore version 2.2.2.6. Branch lengths represent genetic distances. Ultrafast bootstrap and SH-aLRT branch tests were performed with 10,000 replications. Branches with SH-aLRT values ≤ 80% or UFboot values ≤ 95% were labelled with red dots. The leftmost and rightmost panels display zoomed upper and lower parts of the dendrogram, respectively. The cane toad (*Rhinella marina*) labelled with a red arrow.

### Verification of mtDNA genomes from RNASeq data

A near-complete mitochondrial genome with 17,261 to 17,270 bps (95.1% of reference mitochondrial genome) for a total of 132 individuals was generated from the RNASeq and WGS dataset. A sequence with length of 889 bases consisting of repeats in the HV2 section of the control region was excluded due to difficulty in correctly mapping sequencing reads on repeat regions. To determine the accuracy of our RNASeq-derived mtDNA sequences, we then compared the sequences of the six individuals for which we had both RNASeq and WGS data (Table S4). We identified four sites (site 2244, site, 5162, site 7162, site 7920) that showed intra-individual variation across the two datasets in multiple individuals (i.e. potential RNA editing). Two sites in coding regions (site 267, site 477) had intra-individual differences in only one individual. Additionally, one individual had 12 sites in the control region that differed across datasets (Table S4). We replaced the sites with intra-individual variation in multiple individuals with Ns, minimizing variation for subsequent population analyses that might be introduced by RNA editing.

### NUMT analysis

We found 42 mtDNA fragments (38 – 140 bps in length) that mapped to 42 individual contigs of the cane toad draft nuclear genome with over 80% identity (Table S5). All fragments corresponded to the control region of the mitogenome. After the draft genome was masked for repeat elements, no mtDNA fragments were found to map to the nuclear genome.

### Population Genetic Analyses

We identified 262 (1.30%) polymorphic sites in the RNASeq dataset, 198 of which were parsimony-informative, including 25 from RNA genes, 145 from protein-coding genes, and 28 from non-coding regions and the control region (Table S2), which resulted in the identification of 58 haplotypes (Table 1; Figure 4). The thirteen native samples contained twelve unique haplotypes. Two haplotypes found in all introduced populations accounted for nearly 50% of the sampled individuals. Within introduced populations, a “complex-star” shape topology with mostly singleton mutants was observed and 41 of the 46 introduced haplotypes (89%) were private to a single introduction (Hawai’i: 10; Australia: 31).

**Figure 4:**
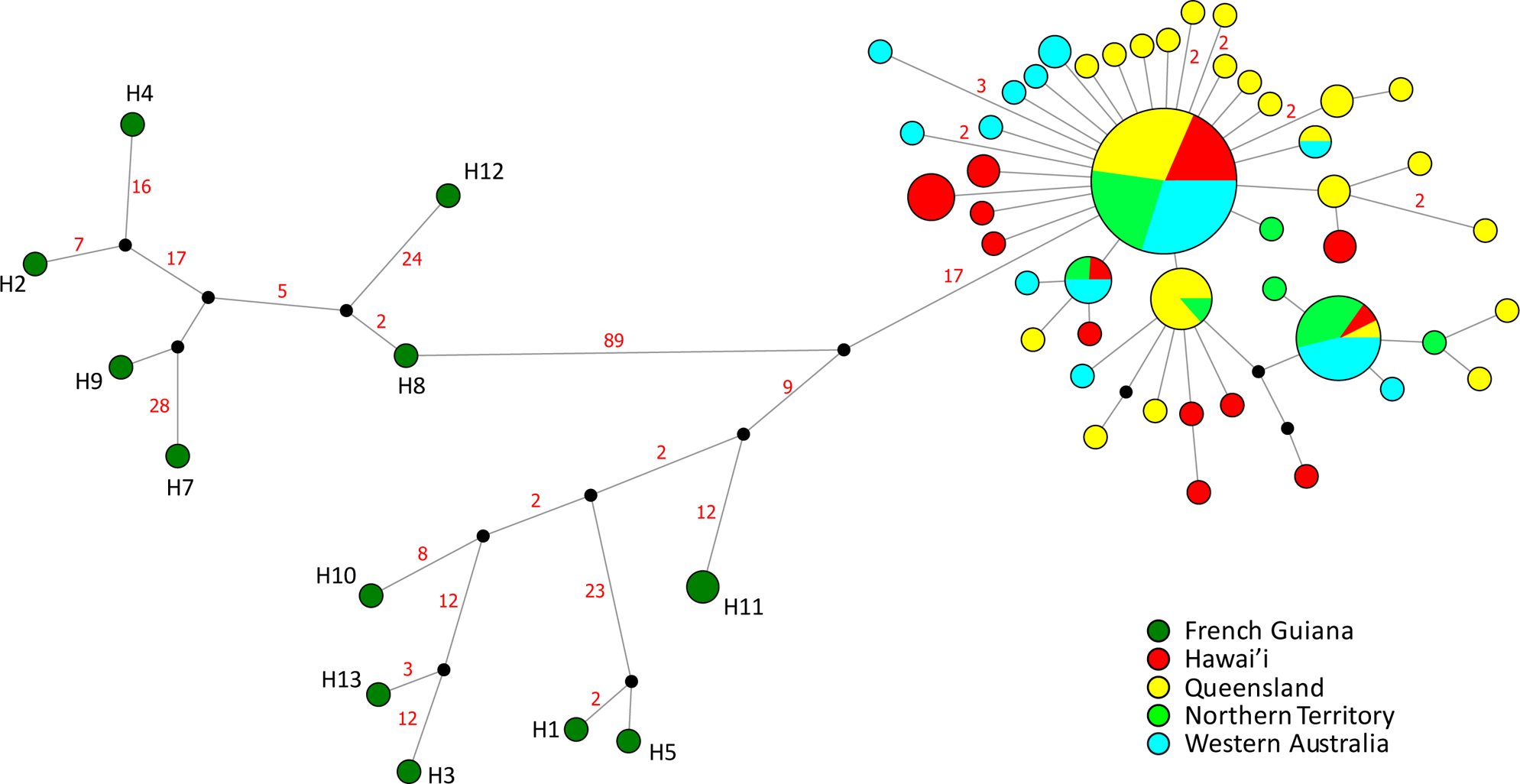
The haplotype network illustrating the relationships among 132 individuals from French Guinea, Hawai’i, and Australia based on near-complete mitochondrial genomes. Each sample location is represented by a different colour. A black solid circle represents a putative haplotype that was not directly sampled. The red number adjacent to a line indicates the number of nucleotide substitutions between haplotypes, while the absence of a number indicates a single nucleotide substitution.

**Table 1:** Population genetics statistics of native range population (French Guiana) and introduced populations (Hawai’i and Australia).

| | *N* | Polymorphic sites | *H* | $\pi \pm SD$ | $h \pm SD$ | HR |
| --- | --- | --- | --- | --- | --- | --- |
| French Guiana | 13 | 205 | 12 | 0.00469 ± 0.00037 | 0.987 ± 0.035 | 12 |
| Hawai'i | 25 | 11 | 13 | 0.00010 ± 0.00002 | 0.880 ± 0.051 | 10.7 |
| Australia | 94 | 40 | 36 | 0.00009 ± 0.00001 | 0.834 ± 0.035 | 12.5 |
| Queensland | 42 | 27 | 23 | 0.00012 ± 0.00001 | 0.890 ± 0.040 | 13.5 |
| Northern Territory | 20 | 5 | 7 | 0.00005 ± 0.00001 | 0.711 ± 0.089 | 6.4 |
| Western Australia | 32 | 16 | 13 | 0.00008 ± 0.00001 | 0.809 ± 0.06 | 9.6 |

*N*: Number of samples; *H*: Number of haplotypes; $\pi$ : nucleotide diversity; *h*: haplotype diversity; HR; haplotype richness.

We also compared the nearly-complete mitochondrial genome dataset with the *ND3* dataset to demonstrate the consistency of haplotype inference across studies, and to put our data into the context of previous work. A total of 24 haplotypes were identified in this analysis (Figure 5); six haplotypes were identified in French Guiana/Guyana, one haplotype was identified in Indonesian samples, three in Hawai’ian and Australian samples, and the remaining haplotypes were from other South America localities. Haplotypes from either side of Andes clearly formed distinct lineages which is consistent to previous work (Acevedo et al., 2016). The single Australian haplotype (Hap1) previously identified in (Slade & Moritz, 1998) was the most common haplotype in our Australian samples and the other two haplotypes (Hap2 and Hap3) within Australia were a single nucleotide different to this common haplotype.

**Figure 5:**
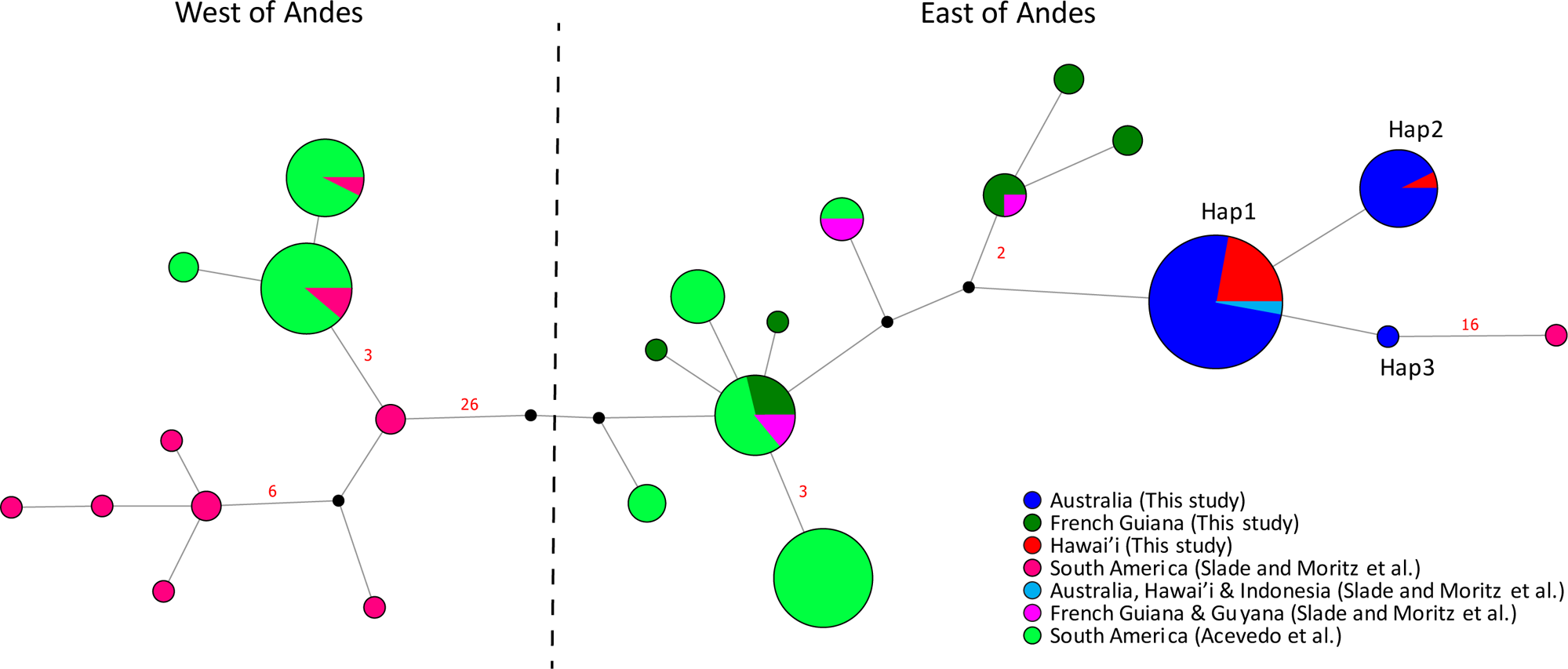
Haplotype network of a 510 bp segment of ND3 with flanking tRNAs, a total of 223 samples, including samples from this study, Slade and Moritz (1998) and Acevedo et al. (2016). Sample locations are represented by different colours. A black solid circle represents a putative haplotype that was not directly sampled. The red number adjacent to a line indicates the number of nucleotide substitutions between haplotypes, while the absence of a number indicates a single nucleotide substitution. Samples are separated by geographical locations on each side of the Andes Mountains in South America (Acevedo et al., 2016). Toads on the west side of the Andes are now known as *Rhinella horribilis*.

Haplotype diversity was highest in the native range, with reduced diversity in both Hawai’ian and Australian populations (*h*: 0.9782 vs 0.8800 and 0.8344, respectively) (Table 1, Figure 6A). Native toads had the highest nucleotide diversity (π: 0.00469) compared to considerably lower diversity in Hawai’ian and Australian toads (π: 0.00010 and 0.00009, respectively) (Table 1, Figure 6B). Queensland toads had similar haplotype diversity but higher nucleotide diversity and haplotype richness to Hawai’ian toads. The population from the Northern Territory had noticeably lower genetic diversity and haplotype richness than other Australian populations (Table 1, Figure 6A, C). When the relationship between haplotypes across the invasion trajectory is considered, the twelve haplotypes from French Guiana were highly dissimilar to those from the Australian and Hawai’ian populations, forming two distinct clades (Figure 4). These were separated from each other by a minimum of 110 base pair changes and from the common invasive haplotype by a minimum of 38 base pair changes (Figure 4). Australian and Hawai’ian haplotypes were intermixed and most of the haplotypes were one base pair different to other haplotypes, with invasive haplotypes being separated from at least one other haplotype by no more than three base pair changes.

**Figure 6:**
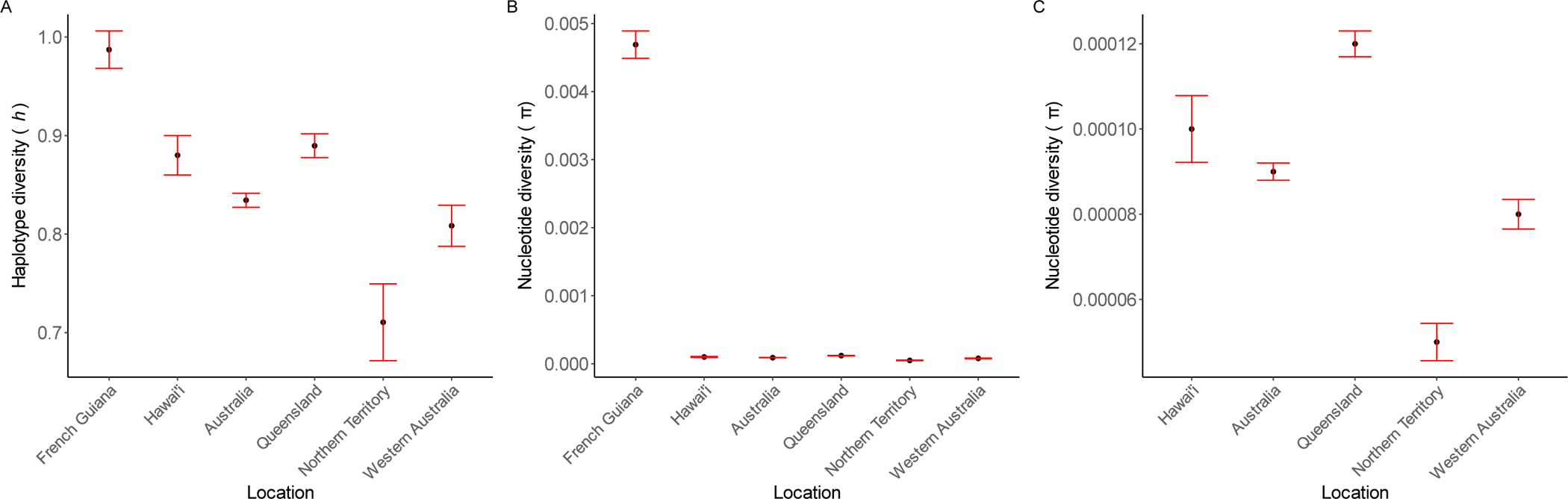
Haplotype diversity (A) and nucleotide diversities (B and C) of native population and introduced populations.

AMOVA results showed that the overall genetic variation among populations (64.49%) was larger than the variation within populations (36.33%) (Table 2). Pairwise *Fst* values were significant between the native population and all introduced populations (*Fst*: 0.55 – 0.67; Table 3). The *Fst* values between introduced populations were all less than 0.1, indicating substantially less differentiation in the invasive range than in the native range. The Hawai’ian population was significantly differentiated from Australian populations (*Fst*: 0.02, *p* = 0.02), but this difference was driven by the Northern Territory (*Fst*: 0.05, *p* = 0.01 and Western Australian populations (*Fst*: 0.03, *p* = 0.01); Queensland was not significantly differentiated to the Hawai’i population (*Fst*: 0.01, *p* = 0.06). Within Australia, the Queensland population showed significant differentiation to both the Northern Territory and Western Australian populations (*Fst*: 0.04, *p* = 0.01 and 0.03, *p* = 0.003, respectively), whereas the Northern Territory and Western Australian populations were not genetically differentiated (*Fst*: −0.01, *p* = 0.72).

**Table 2:** Analysis of molecular variance (AMOVA) of all populations.

| Source of variation | *df* | Sum of squares | % of variation | Fixation indices |
| --- | --- | --- | --- | --- |
| All populations |  |  |  |  |
| Among groups | 2 | 482.96 | 64.49 |  |
| Among populations<br>within groups | 2 | 2.91 | -0.82 |  |
| Within populations | 127 | 579.35 | 36.33 | $F_{ST} = 0.64^{***}$ |

\*\*\*: $p < 0.001$

**Table 3:**
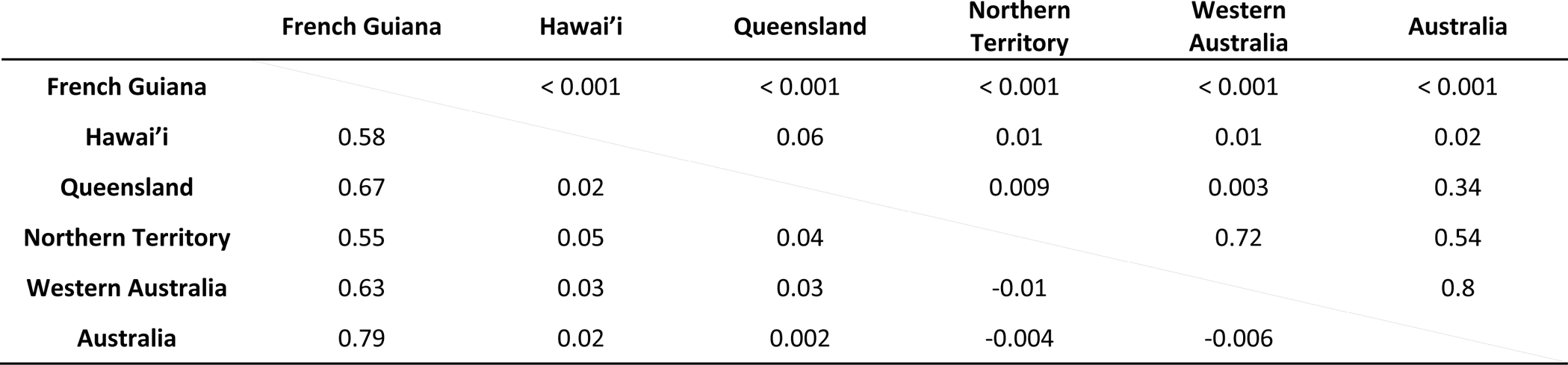
Population differentiation (pairwise *Fst*) between native range and introduced ranges (Hawai’i and Australia) (below the diagonal line) The associated *p*-values were shown above the diagonal line.

|  | French Guiana | Hawai'i | Queensland | Northern Territory | Western Australia | Australia |
| --- | --- | --- | --- | --- | --- | --- |
| French Guiana |  | < 0.001 | < 0.001 | < 0.001 | < 0.001 | < 0.001 |
| Hawai'i | 0.58 |  | 0.06 | 0.01 | 0.01 | 0.02 |
| Queensland | 0.67 | 0.02 |  | 0.009 | 0.003 | 0.34 |
| Northern Territory | 0.55 | 0.05 | 0.04 |  | 0.72 | 0.54 |
| Western Australia | 0.63 | 0.03 | 0.03 | -0.01 |  | 0.8 |
| Australia | 0.79 | 0.02 | 0.002 | -0.004 | -0.006 |  |

Two neutrality tests were implemented to study the demographic history of cane toad populations: Fu’s *Fs* and Ramos-Onsins and Rozas’ *R_2_*statistic (Table 4). The native population had non-significant positive Fu’s *Fs* values and a large *R_2_* statistic, indicating the population is not expanding or shrinking (i.e. is demographically stable). Fu’s *Fs* values were negative in all introduced populations, indicating an excess of rare haplotypes compared to the expectation from neutrality and all these were statistically significant. *R_2_* statistics were significant in Hawai’i, Queensland and Western Australia’s populations.

**Table 4:** Demographic estimators of native range population (French Guiana) and introduced populations (Hawai’i and Australia).

| | Fu's $F_s$ | $R_2$ | Demographic expansion | | Spatial expansion | |
| --- | --- | --- | --- | --- | --- | --- |
|  |  |  | SSD | Raggedness | SSD | Raggedness |
| **French Guiana** | 1.29<br>($p = 0.66$) | 0.18<br>($p = 0.891$) | 0.03<br>($p = 0.19$) | 0.03<br>($p = 0.39$) | 0.03<br>($p = 0.08$) | 0.03<br>($p = 0.55$) |
| **Hawai'i** | -8.78<br>($p < 0.001$) | 0.08<br>($p = 0.02$) | 0.01<br>($p = 0.44$) | 0.080<br>($p = 0.24$) | 0.01<br>($p = 0.31$) | 0.08<br>($p = 0.21$) |
| **Australia** | -27.92<br>($p < 0.001$) | 0.02<br>($p < 0.001$) | 0.002<br>($p = 0.30$) | 0.067<br>($p = 0.16$) | 0.002<br>($p = 0.20$) | 0.07<br>($p = 0.14$) |
| **Queensland** | -23.04<br>($p < 0.001$) | 0.03<br>($p < 0.001$) | 0.001<br>($p = 0.70$) | 0.05<br>($p = 0.38$) | 0.002<br>($p = 0.66$) | 0.05<br>($p = 0.42$) |
| **Northern Territory** | -3.91<br>($p < 0.001$) | 0.10<br>($p = 0.072$) | 0.03<br>($p = 0.07$) | 0.21<br>($p = 0.03$) | 0.03<br>($p = 0.03$) | 0.21<br>($p = 0.05$) |
| **Western Australia** | -8.950<br>($p < 0.001$) | 0.05<br>($p = 0.001$) | 0.003<br>($p = 0.58$) | 0.08<br>($p = 0.34$) | 0.003<br>($p = 0.35$) | 0.08<br>($p = 0.26$) |

Mismatch distribution graphs of all populations are shown in Figure S1. The native population showed a multimodal distribution with non-significant SSD and raggedness values, indicating support for spatial and demographic expansion (Table 4). Most introduced populations showed a unimodal distribution for both expansion models with non-significant SSD and raggedness indices (Table 4), supporting the presence of spatial and demographic expansions. However, the Northern Territory population had significant or marginally significant p-values for SSD and raggedness values for both expansion models indicating poor support (spatial expansion: SSD *p*-value = 0.03, raggedness *p*-value = 0.05; demographic expansion: SSD *p*-value = 0.07, raggedness *p*-value = 0.03).

## Discussion

This study uses mitochondrial genome sequences to study the evolutionary and phylogenetic relationships of the iconic invasive cane toad. Here, we constructed the first reference mitochondrial genome of *R. marina* and placed this species within the order Anura. We utilised RNASeq-derived mitogenomes to illustrate a greater number and diversity of haplotypes in mtDNA present in Australia and Hawai’i, in contrast to the surprisingly low genetic diversity previously described using the *ND3* gene alone. Our conclusions are further supported by evidence of demographic and spatial expansion in invasive cane toad populations. This study provides new insights into the diversity of native cane toads and how they have evolved along their invasion trajectory.

### Phylogenetic inference

The phylogenetic relationships of toads within Anura are understudied. More specifically, the cryptic *Rhinella* species complex remains poorly understood (Pereyra et al., 2021). Zhang et al. (2013) attempted to study the Anuran phylogeny using partial mitochondrial genomes. However, no *Rhinella* species were included in their study. Currently, the evolutionary relationship of *Rhinella* species in the order Anura has yet to be fully documented using complete mitochondrial genome or even nuclear genes and/or mitochondrial genes. To address this knowledge gap, we generated the first complete mitochondrial genome from any *Rhinella* species and investigated the phylogenetic relationship of *Rhinella* to other Anurans. Although the topology of our tree confirmed previous hypotheses (e.g., in line with Feng et al. (2017), *Rhinella* formed a sister clade with *Anaxyrus*), it also yielded different conclusions in some aspects. For example, Feng et al. (2017) built a tree for frog species using 95 unlinked nuclear genes and suggested that the genus *Bufo* is monophyletic with respect to *Duttaphrynus*. In contrast, our whole-mitochondrial genome phylogenetic analysis found that the genus *Bufo* is a subclade of *Duttaphrynus*. Our study provides one of the most comprehensive phylogenetic analyses of the relationships within the order Anura to date and provides a starting point for future phylogenetic analyses within the genus *Rhinella*.

Choosing to use a complete dataset of a few selected loci versus a large supermatrix with missing data for phylogenetic inference remains a controversial issue (see for review (Lemmon, Brown, Stanger-Hall, & Lemmon, 2009; Roure, Baurain, & Philippe, 2013; Wiens & Morrill, 2011)). Typically, phylogenetic trees are generated using single or multiple nuclear genes, single or multiple nuclear genes plus mitochondrial genes, or multiple nuclear genes with morphological trait matrices, but the use of complete mitochondrial genomes for phylogenetic inference is rare due to resource limitations. Missing data in phylogenetic analyses can bias phylogenetic calculation and tree branch lengths (Lemmon et al., 2009; Xia, 2014). Nonetheless, contrasting findings have also suggested that when a sufficient number of loci and taxa are included, phylogenetic signals remain consistent regardless of whether or not the dataset includes missing data (Molloy & Warnow, 2018; Shavit Grievink, Penny, & Holland, 2013). Despite using complete mitochondrial genomes in this study, 11% of branches remained statistically unsupported, and the genera *Bufo*, *Bufotes*, *Anaxyrus* and *Rhinella* in the Bufonidae family are also not well-supported. This result suggests that the resolution power of the complete mitochondrial genome may not be sufficient, and/or more representative sampling of different anuran species may be needed to provide more robust support for statistical calculations and improve our understanding of the phylogeny.

### Comparison between RNASeq & WGS results

The use of transcriptomic data to build mitochondrial genomes has been suggested as a cost-effective alternative to long PCRs or direct sequencing of total DNA (Forni et al., 2019), but it is important to ensure that the inferred polymorphisms accurately reflect genetic variation. In our study, comparing mtDNA data obtained from WGS and RNASeq, we found discrepancies in multiple nucleotide positions, all of which were in rRNA, tRNA and the control region, which could be the result of post-transcriptional modifications or RNA editing which are well-recognised phenomena across multiple taxa (Levanon et al., 2004; Porath, Knisbacher, Eisenberg, & Levanon, 2017; St Laurent et al., 2013). While common substitutions such as adenosine to inosine (A-to-I) deamination and cytidine to uridine (C-to-U) pyrimidine exchange were found, we also observed less common substitutions (e.g. thymine to cytidine (T-to-C) that have been previously documented in another toad species, *Xenopus tropicalis* (Zaranek, Levanon, Zecharia, Clegg, & Church, 2010). These processes are important for creating multiple transcript isoforms and gene regulation (Tang, Fei, & Page, 2012) and are known to be associated with adaptation (Duan, Dou, Luo, Zhang, & Lu, 2017; Duan et al., 2021). RNA editing of the tRNA anticodon recognition site would extend the codon matching ability and further increase the variation of the resulting translated proteins. Although we did not observe nucleotide differences in position 34 or position 37 of tRNA, which are typically recognised for deamination of adenosine to inosine which can be paired with A, C or U residues (Jackman & Alfonzo, 2013), we did find changes in the D arm of tRNA (Aspartic acid and Lysine), the T arm of tRNA (Phenylalanine) and the acceptor stem of tRNA (Tryptophan) (Figure S2), which are critical for the recognition of aminoacyl tRNA synthetases to bind appropriate amino acids to its designated tRNA (Ganesh & Maerkl, 2022). Failure to form a tRNA-amino acid complex could inhibit the use of a specific amino acid in subsequent protein translation. Further proteome analysis is required to determine if these changes affect the protein translation machinery.

Post-transcriptional modifications were more diverse in the native populations than in the invasive populations. In particular, one individual from French Guiana displayed discrepancies in the control region between WGS and RNASeq data, all of which were transitions. This observation might result from tissue-specific expression in spleen or RNA editing. Studies of RNA editing in mtDNA have primarily focused on protein-coding genes and tRNAs, with less research conducted on the control region of the mtDNA, which is responsible for regulating RNA and DNA synthesis. While the implications of potential RNA editing in cane toads remain unclear, these events may contribute to diversity at a transcriptional and translational level. The low diversity of post-transcriptional modification in the invasive populations might be the result of selective sweep for favourable haplotypes with low activity of RNA-editing enzymes.

NUMTs, or nuclear insertions of mitochondrial DNA fragments (Lopez, Yuhki, Masuda, Modi, & O’Brien, 1994), can potentially present another source of genetic variation and contribute to false-positive signals of variation in nuclear data (Song, Buhay, Whiting, & Crandall, 2008). NUMTs have been reported in many different taxa including vertebrates, plants and protists (Hazkani-Covo, Zeller, & Martin, 2010). They vary in size, ranging from minimal insertions to hundreds of kilobases and tend to be longer in species with larger nuclear genomes (Hazkani-Covo et al., 2010; Ko & Kim, 2016). Despite NUMTs generally being perceived as non-functional due to their ability to shift reading frames, the presence of in-frame stop codons and the use of different genetic codes, the transcription of NUMTs into non-coding RNA molecules can be captured by RNASeq and potentially falsely interpreted as mitochondrial variation among or within individuals. NUMTs studies on amphibian species are limited. In *Xenopus tropicalis*, over 10kb of NUMTs has been found in the nuclear genome (Hazkani-Covo et al., 2010). Investigation of the presence/absence of NUMTs in cane toad nuclear genome and the potential expression of NUMTs is required to distinguish between genuine mtDNA variation within individuals and false-positive signals caused by incorporating nuclear genetic data into mitochondrial genetic analyses.

A search for NUMTs in the published cane toad draft genome (Edwards et al., 2018) yielded only matches with simple repeat regions within the mtDNA control region. In other species, NUMTs originate across the entire mtDNA genome (Dayama, Zhou, Prado-Martinez, Marques-Bonet, & Mills, 2020; Wei et al., 2022). Following the repeat-masking of the draft nuclear genome, we found no evidence of NUMTs suggesting that the putative NUMTs were just low-complexity regions in the nuclear genome. Similar to other gene-searching methods such as BUSCO, the accuracy of the search algorithm heavily depends on the contiguity of the genome and the divergence of the query from the database subjects. Given the high number of contigs in the cane toad draft genome, potential NUMTs hits may have been missed through breakpoints. Therefore, it is important to develop a cane toad genome assembly with higher contiguity to improve our ability to investigate NUMTs. Additionally, ancient NUMTs may be too divergent to be detected by the methods employed, but NUMTs with such low identity would not interfere with mtDNA analysis.

It is worth noting that the consensus sequences generated from the bioinformatic pipelines used in this study represented the most common nucleotide at positions where multiple nucleotides were found. Although these discrepancies could result from sequencing errors, our filtering is likely to have prevented this. As discussed above, RNA editing or NUMTs could have resulted in putative polymorphisms within individuals in our RNASeq dataset. It is also possible that in some cases, heteroplasmy could have caused these putative polymorphisms. While determining the cause of this phenomenon is beyond the scope of this study, it is unlikely that this has affected population-level statistics.

### Population genetic analyses

Interestingly, both the haplotype network (Figure 4) and the multimodal mismatch distribution (Figure S1) indicate the presence of multiple genetically divergent lineages within native populations. Notably, individuals within each lineage are intermixed, rather than being grouped by geographical locations. For instance, despite an 80 km geographical separation, haplotypes H4 and H7 are found in the same lineage while haplotypes H12 and H11 were collected only 9 m apart but belong to different lineages. Native-range cane toads can travel 60-180 m per move for sheltering and foraging purposes (Shine et al., 2021). However, the overall dispersal rates are quite low, as the toads ultimately tend to return to the same shelters (DeVore, Shine, & Ducatez, 2021). Consequently, the possibility of hybridisation between toads from these distinct lineages exists. However, a recent study in South and Central America which utilised ddRAD sequencing has offered a different perspective on the phylogenetic relationship of cane toads. This study revealed toads from the coastal area formed two sister clades, distinct from the clade formed by toads from the rainforest regions (Mittan-Moreau et al., 2022). It becomes crucial to ascertain the precise populations of cane toads from the native range that were used as founding individuals, especially given the limited comprehensive description in prior research (Easteal, 1981). In Shine et al. (2021), cane toads from the range-edge regions (Northern Territory and Western Australia) exhibited similar dispersal patterns to native toads in one of the coastal sites. This suggests the adaptation of cane toads in hot and arid environments is likely facilitated by the standing genetic variation(s) inherited from the progenitors utilising dry, saline coastal beach environments (DeVore et al., 2021), rather than relying solely on the genetic adaptation and selection of novel mutation(s). A more thorough investigation into these distinct lineages becomes imperative for unravelling the mechanisms underlying the adaptations of cane toads in Australia.

Our study challenges the assumption of low genetic diversity in Hawai’ian and Australian populations. Slade and Moritz (1998) reported a unique haplotype at *ND3* within Australian populations, in comparison with multiple haplotypes in native populations in Central and South America. This result supported the presence of founder effects following the introduction of cane toads into Hawai’i and Australia. Here, we revealed two additional, previously undocumented, *ND3* haplotypes (Hap2 and Hap3) in Hawai’ian and Australian populations, each of which differed by only one nucleotide from the previously identified haplotype (Hap1) (Figure 5). In comparison to our analysis using nearly complete mitogenomes discussed above, this analysis highlights that a single gene is not sufficient to accurately infer genetic diversity.

The significant decrease in haplotype diversity and nucleotide diversity observed in both Hawai’ian and Australian populations, compared to native populations, does support that genetic bottlenecks and founder effects occurred post-introduction (Table 1, Figure 6). This finding is consistent with previous studies on this species using nuclear genetic markers including SNPs, microsatellite markers, and MHC class I and II loci (Lillie et al., 2016; Lillie et al., 2014; Selechnik, Richardson, Shine, DeVore, et al., 2019). In this study, all three neutrality indices showed strong signals of population expansion and the observed data fitted to both demographic expansion and spatial expansion models in most invasive sampling sites (Table 4). This conclusion is supported by the star-like shaped topology of the haplotype network created using the nearly-complete mitochondrial genome dataset. Despite reduced diversity in the introduced range, the degree of the decrease is less than previously reported (Leblois et al., 2000). Additionally, the star-shaped topology of the network containing the introduced haplotypes, the absence of native samples here, and the significant negative Fu’s *Fs* values indicate that many of the singleton haplotypes discovered in Hawai’i and Australia may be the products of post-introduction mutations. Alternatively, due to the limited sample sizes in the native range, it is possible that any introduced haplotypes exist but were not included in our sample set for analysis.

We observed a curvilinear pattern of genetic diversity indices across Australian populations (from Queensland to Western Australia), a pattern also found in spleen gene expression (Selechnik, Richardson, Shine, Brown, et al., 2019), limb length (Hudson, McCurry, Lundgren, McHenry, & Shine, 2016), spleen and body mass (Gregory P. Brown, Kelehear, Shilton, Phillips, & Shine, 2015). The mechanism behind this reduction of genetic diversity in Northern Territory toads remains unclear, but it may be due to demographic processes that reduce the effective population size and increase genetic drift (Nei & Tajima, 1981), genetic hitchhiking that removes neutral polymorphic sites under positive selection (Fay & Wu, 2000) or directional selection due to harsh climate (Selechnik, Richardson, Shine, DeVore, et al., 2019), possibly in combination with dispersion-driven spatial sorting at the invasion front (Clarke, Shine, & Phillips, 2019). Despite our finding that the Northern Territory and Western Australia populations were not genetically differentiated, it is likely that they have experienced different local evolutionary forces and demographic processes. Northern Territory toads, unlike other populations, only exhibited demographic expansion and not spatial expansion, as supported by lower levels in all statistical measures.

Our results demonstrate significant differentiation between native and introduced populations (Table 3). We identified three genetic clusters: 1) native toads, 2) Hawai’i and Queensland toads (*Fst*: 0.016, *p* = 0.056) and 3) Northern Territory and Western Australia toads (*Fst*: −0.012, *p* = 0.72), consistent with previous results using SNPs from the nuclear genome (Selechnik, Richardson, Shine, DeVore, et al., 2019). It is interesting that the Queensland toads are not differentiated from those sampled in Hawai’i but differ from toads sampled in the Northern Territory and Western Australia. Queensland toads have been separated from their Hawai’ian conspecifics for almost 90 years but do share similar environmental conditions (Selechnik, Richardson, Shine, DeVore, et al., 2019). Northern Territory and Western Australian toads suffer from reduced genetic diversity compared to Queensland toads, so they have likely experienced more genetic drift, which may explain this difference. However, it is also possible that the striking reduction in precipitation and increase in temperature experienced by Northern Territory and Western Australian toads compared to those from Queensland has resulted in selective differences in the mitogenome between these populations. In support of this possibility, significant negative Fu’s *Fs* can indicate the presence of selection.

## Conclusion

Our study used the whole mtDNA genome as a genetic marker to investigate the evolution of cane toads across their invasion trajectory from the native range to Australia, adding to our knowledge of changes to genetic diversity as cane toads have been serially introduced to multiple locations. Unsurprisingly, we found reduced genetic diversity in introduced populations, suggesting that genetic bottlenecks have occurred in these populations. Despite this reduction, our data indicate greater genetic diversity in introduced populations than has been previously described using mtDNA. We identified signatures of demographic and spatial expansion in most introduced localities we sampled, consistent with ongoing invasion. We described 46 haplotypes in introduced populations, mostly consisting of single base-pair changes, suggesting evolution following introduction. By the comparison of WGS- and RNASeq-derived mitogenomes, we have identified the presence of post-transcriptional modification or RNA editing, which may supply additional genetic diversity upon which selection can act.

## Acknowledgement

We thank Mark Richardson and Yini Peng Lee for assistance with genomic sequencing and processing of genomic data and the Ramaciotti Centre for Genomics, University of New South Wales (UNSW) and Deakin Genomic Centre, Deakin University who performed the sequencing in this study. This research includes computations using the computational cluster Katana supported by Research Technology Services at UNSW Sydney. This study was supported by the Australian Research Council (grants DP160102991, DP190100507, to RS and LAR), the UNSW Scientia Program (to LAR), and the UNSW Scientia PhD Scholarship (to KC).

## Author Contributions

KC, RS, RJE, and LAR developed this project. JLD and SD conducted field work. LAR conducted DNA extractions. KC, TGA, RJE, and LAR analysed the data. All authors contributed to data interpretation and writing.

## Data availability

The haplotype sequences will be available on the NCBI public database once the manuscript is accepted.

