## Supplementary Information for "Mitochondrial genome analyses uncover intriguing population structure across the invasive trajectory of an iconic invader"

Figure S1: Mismatch distribution graphs of native and introduced populations. The x-axis represents the number of pairwise nucleotide differences between pairs of samples. The y-axis represents the frequency of the comparisons. Both the demographic expansion model (solid red line) and the spatial expansion model (solid blue line) were investigated.

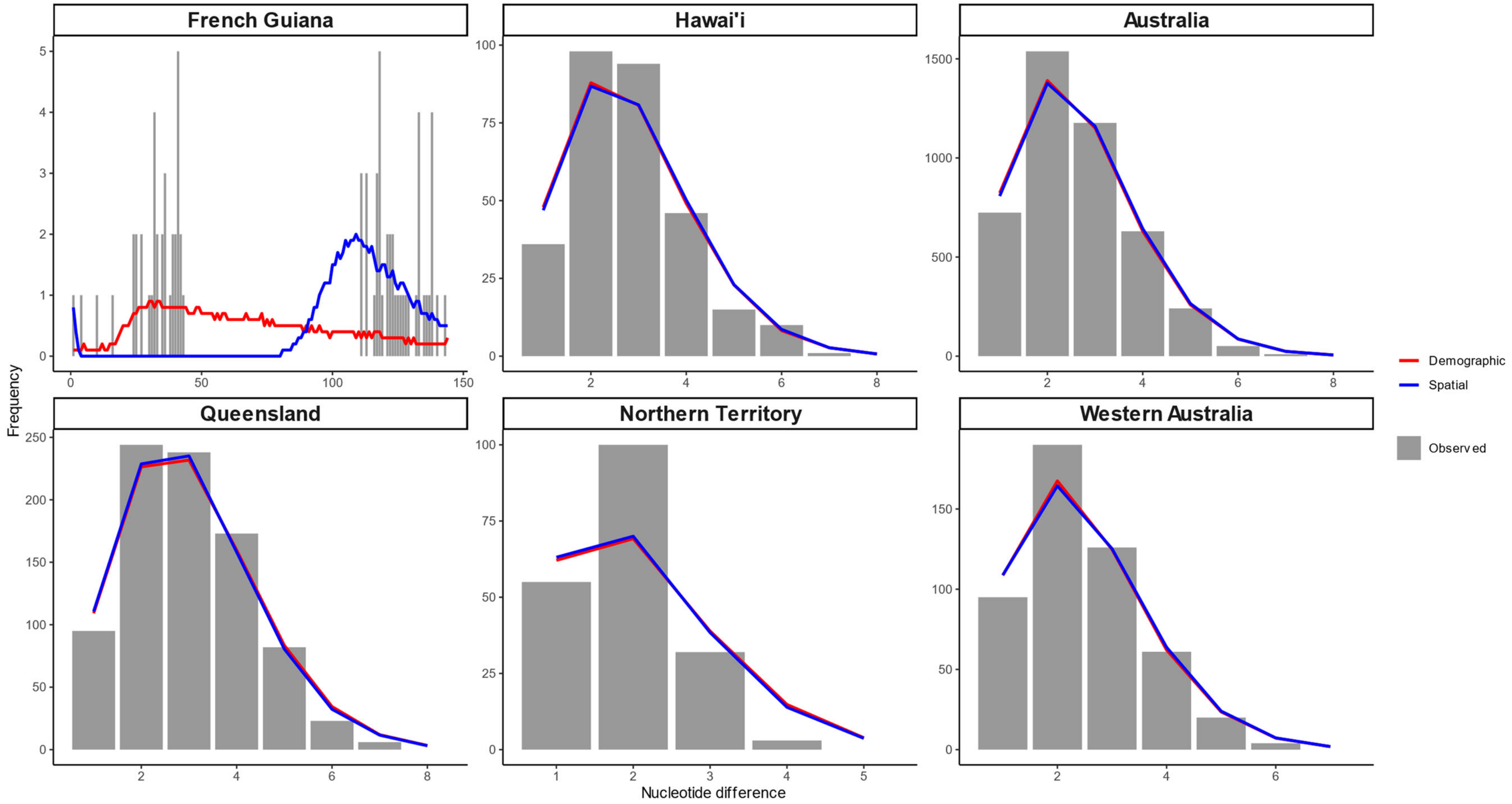

Figure S2: Predicted tRNA secondary structures showing nucleotide substitutions between WGS and RNASeq data. Nucleotides marked with red circles indicate differences between the WGS and RNASeq datasets. (a) tRNA (Lysine), (b) tRNA (Aspartic acid), (c) tRNA (Tryptophan), and (d) tRNA (Phenylalanine) from the WGS dataset (left) and the RNASeq dataset (right).

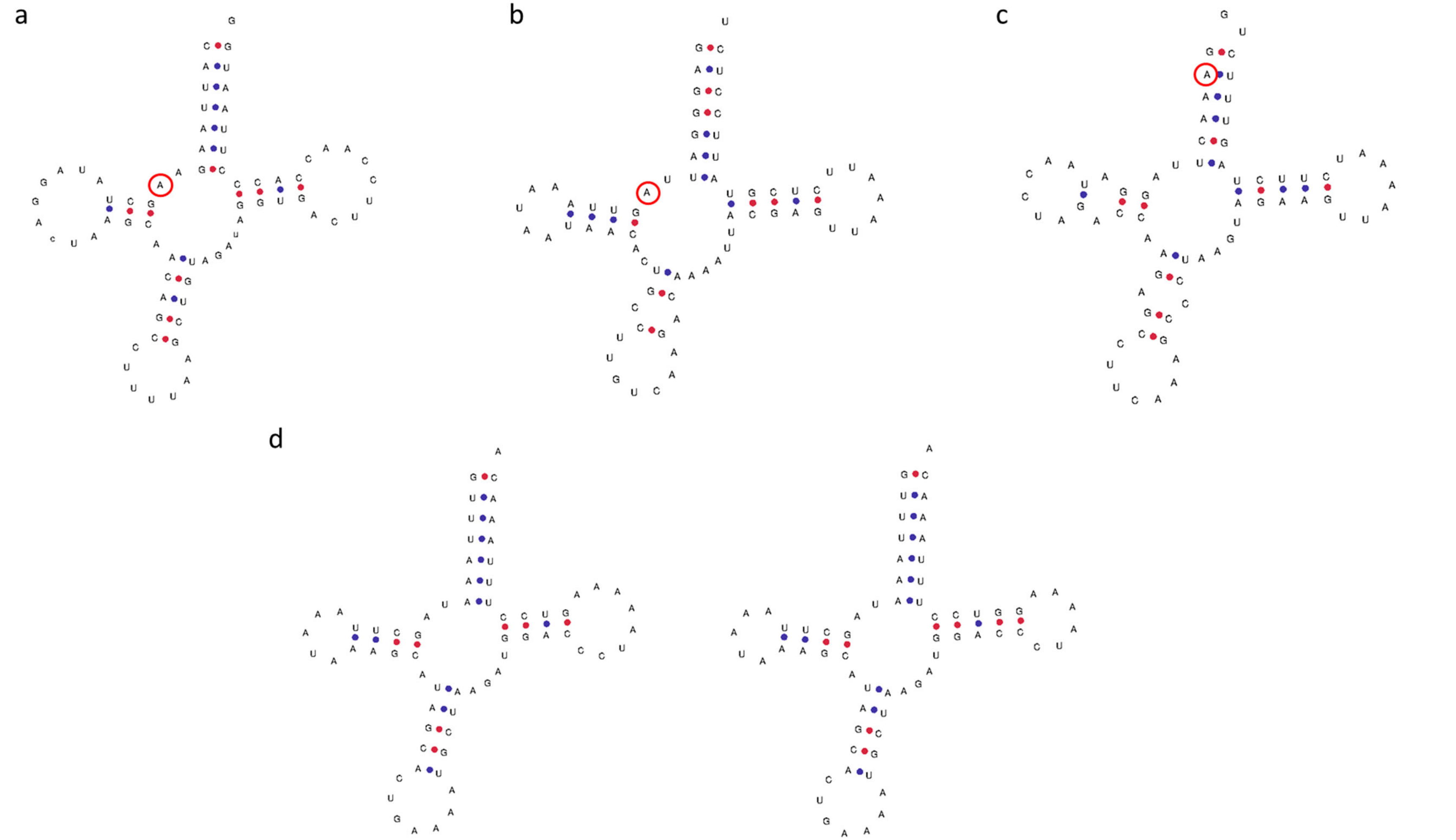

**Table S1:** ***Rhinella marina*** **samples used in this study.**

| Site | Location | Sample ID | Type of sequencing | Haplotype notation | Latitude | Longitude | Reference |
|---|---|---|---|---|---|---|---|
| **French Guiana** | | | | | | | |
| | Plage de Montjoly | RMF010 | RNA | H7 | 4.913 | -52.261 | |
| | Plage de Montjoly | RMF022 | RNA | H9 | 4.913 | -52.267 | |
| | Plage de Montjoly | RMF044 | WGS | H3 | 4.913 | -52.260 | |
| | Plage de Montjoly | RMF046 | RNA | H11 | 4.913 | -52.259 | |
| | Plage de Montjoly | RMF047 | RNA | H12 | 4.913 | -52.259 | |
| | Plage de Montjoly | RMF048 | RNA, WGS | H11 | 4.913 | -52.259 | |
| | Plage de Montjoly | RMF049 | RNA | H13 | 4.913 | -52.260 | This study |
| | Plage de Gosselin | RMF017 | RNA | H8 | 4.891 | -52.253 | |
| | Plage de Gosselin | RMF031 | WGS | H1 | 4.890 | -52.252 | |
| | Route des Plages Drain | RMF028 | RNA | H10 | 4.879 | -52.246 | |
| | Past Regina wetland | RMF042 | WGS | H2 | 4.203 | -52.126 | |
| | Past Regina wetland | RMFG0403 | WGS | H4 | 4.203 | -52.126 | |
| | Regina Wash | RMFG107 | WGS | H5 | 4.363 | -52.280 | |
| | Lake Maracaibo, Venezuela | | | | | | |
| | Higuerote, Venezuela | | | | | | |
| | Guiria, Venezuela | | | | | | |
| | Cayenne, French Guiana | | | | | | |
| | Madre de Dios, Atalaya, Peru | | | | | | Slade and Moritz et al. |
| | Mexico | | | | | | |
| | Costa Rica | | | | | | |
| | Panama | | | | | | |
| | Texas, USA | | | | | | |
| **Hawai'ian Islands** | | | | | | | |
| O'ahu | | | | | | | |
| | Haiku Gardens | RMH004 | WGS | H6 | 21.415 | -157.816 | |
| | Haiku Gardens | RMH006 | RNA, WGS | H14 | 21.415 | -157.816 | |
| | Haiku Gardens | RMH008 | RNA | H15 | 21.415 | -157.816 | |
| | Haiku Gardens | RMH043 | RNA | H20 | 21.415 | -157.816 | |
| | Haiku Gardens | RMH044 | RNA | H14 | 21.415 | -157.816 | |
| | Haiku Gardens | RMH046 | RNA | H6 | 21.415 | -157.816 | |
| | Haiku Gardens | RMH047 | RNA | H14 | 21.415 | -157.816 | |
| | Haiku Gardens | RMH049 | RNA | H21 | 21.415 | -157.816 | |
| | Haiku Gardens | RMH050 | RNA | H14 | 21.415 | -157.816 | This study |
| | Kapolei Regional Park | RMH018 | RNA | H6 | 21.336 | -158.078 | |
| | Kapolei Regional Park | RMH020 | RNA | H6 | 21.336 | -158.078 | |
| | Kapolei Regional Park | RMH021 | RNA | H16 | 21.336 | -158.078 | |
| | Kapolei Regional Park | RMH024 | RNA | H17 | 21.336 | -158.078 | |
| | Kapolei Regional Park | RMH025 | RNA | H18 | 21.336 | -158.078 | |
| | Kapolei Regional Park | RMH026 | RNA | H6 | 21.336 | -158.078 | |
| | Kapolei Regional Park | RMH027 | RNA | H6 | 21.336 | -158.078 | |
| | Kapolei Regional Park | RMH028 | RNA | H19 | 21.336 | -158.078 | |
| | Manoa | | | | | | Slade and Moritz et al. |
| Hawai'i | | | | | | | |
| | Paradise Park | RMH051 | RNA, WGS | H23 | 19.572 | -154.958 | |
| | Paradise Park | RMH052 | RNA, WGS | H6 | 19.572 | -154.958 | |
| | Paradise Park | RMH059 | RNA | H22 | 19.572 | -154.958 | |
| | Paradise Park | RMH061 | RNA | H22 | 19.572 | -154.958 | This study |
| | Kingslake | RMH075 | RNA | H24 | 19.939 | -155.875 | |
| | Kingslake | RMH077 | RNA | H6 | 19.939 | -155.875 | |
| | Kingslake | RMH079 | RNA | H24 | 19.939 | -155.875 | |
| | Kingslake | RMH081 | RNA | H25 | 19.939 | -155.875 | |
| | Coconut Island | | | | | | Slade and Moritz et al. |
| **Australia** | | | | | | | |
| Western Australia | | | | | | | |
| | Caroline Pool | RM640 | RNA | H21 | -18.227 | 127.760 | |
| | Caroline Pool | RM642 | RNA | H6 | -18.227 | 127.760 | |
| | Caroline Pool | RM645 | RNA | H21 | -18.227 | 127.760 | |
| | Caroline Pool | RM647 | RNA | H21 | -18.227 | 127.760 | |
| | Caroline Pool | RM650 | RNA | H6 | -18.227 | 127.760 | |
| | Durack River | RM657 | RNA | H52 | -15.640 | 127.234 | |
| | Durack River | RM658 | RNA | H53 | -15.640 | 127.234 | |
| | Durack River | RM670 | RNA | H6 | -15.640 | 127.234 | |
| | Durack River | RM676 | RNA | H54 | -15.640 | 127.234 | |
| | Durack River | RM684 | RNA | H21 | -15.640 | 127.234 | |
| | Emma Gorge | RM010 | RNA, WGS | H33 | -15.906 | 128.128 | |
| | Emma Gorge | RM021 | RNA | H6 | -15.906 | 128.128 | |
| | Emma Gorge | RM034 | RNA | H6 | -15.906 | 128.128 | |
| | Emma Gorge | RM039 | RNA | H50 | -15.906 | 128.128 | |
| | Emma Gorge | RM045 | RNA, WGS | H51 | -15.906 | 128.128 | |
| | Purnululu | RM061 | RNA | H6 | -17.430 | 128.304 | This study |
| | Purnululu | RM082 | RNA | H6 | -17.430 | 128.304 | |
| | Purnululu | RM087 | RNA | H21 | -17.430 | 128.304 | |
| | Purnululu | RM092 | RNA | H6 | -17.430 | 128.304 | |
| | Purnululu | RM094 | RNA | H6 | -17.430 | 128.304 | |
| | Purnululu | RMP028 | RNA | H16 | -17.448 | 128.547 | |
| | Purnululu | RMP029 | RNA | H6 | -17.448 | 128.547 | |
| | Purnululu | RMP031 | RNA | H55 | -17.448 | 128.547 | |
| | Purnululu | RMP032 | RNA | H56 | -17.448 | 128.547 | |
| | Purnululu | RMP036 | RNA | H6 | -17.448 | 128.547 | |
| | Purnululu | RMP037 | RNA | H56 | -17.448 | 128.547 | |
| | Purnululu | RMP038 | RNA | H21 | -17.448 | 128.547 | |
| | Purnululu | RMP039 | RNA | H16 | -17.448 | 128.547 | |
| | Purnululu | RMP040 | RNA | H57 | -17.448 | 128.547 | |
| | Purnululu | RMP042 | RNA | H6 | -17.448 | 128.547 | |
| | Purnululu | RMP045 | RNA | H58 | -17.448 | 128.547 | |
| | Purnululu | RMP046 | RNA | H6 | -17.448 | 128.547 | |
| Northern Territory | | | | | | | |
| | Cape Crawford | RM517 | RNA | H6 | -16.667 | 135.800 | |
| | Cape Crawford | RM521 | RNA | H21 | -16.667 | 135.800 | |
| | Cape Crawford | RM523 | RNA | H6 | -16.667 | 135.800 | |
| | Cape Crawford | RM528 | RNA | H48 | -16.667 | 135.800 | |
| | Cape Crawford | RM539 | RNA | H6 | -16.667 | 135.800 | |
| | Cape Crawford | RM542 | RNA | H6 | -16.667 | 135.800 | |
| | Cape Crawford | RM544 | RNA | H49 | -16.667 | 135.800 | |
| | Cape Crawford | RM548 | RNA | H6 | -16.667 | 135.800 | |

| | | | | | | | |
|---|---|---|---|---|---|---|---|
| | Mataranka | RM510 | RNA | H6 | -14.923 | 133.066 | |
| | Mataranka | RM511 | RNA | H47 | -14.923 | 133.066 | This study |
| | Mataranka | RM515 | RNA | H6 | -14.923 | 133.066 | |
| | Mataranka | RM581 | RNA | H21 | -14.923 | 133.066 | |
| | Timber Creek | RM604 | RNA | H21 | -15.643 | 130.467 | |
| | Timber Creek | RM607 | RNA | H21 | -15.643 | 130.467 | |
| | Timber Creek | RM608 | RNA | H21 | -15.643 | 130.467 | |
| | Timber Creek | RM704 | RNA | H6 | -15.643 | 130.467 | |
| | Timber Creek | RM712 | RNA | H37 | -15.643 | 130.467 | |
| | Timber Creek | RM713 | RNA | H16 | -15.643 | 130.467 | |
| | Timber Creek | RM714 | RNA | H6 | -15.643 | 130.467 | |
| | Timber Creek | RM726 | RNA | H6 | -15.643 | 130.467 | |
| | Borroloola | | | | | | Slade and Moritz et al. |
| Queensland | | | | | | | |
| | Burketown | RM349 | RNA | H6 | -17.852 | 139.633 | |
| | Burketown | RM350 | RNA | H6 | -17.852 | 139.633 | |
| | Burketown | RM352 | RNA | H39 | -17.852 | 139.633 | |
| | Burketown | RM354 | RNA | H40 | -17.852 | 139.633 | |
| | Cairns | RMP001 | RNA | H43 | -16.919 | 145.778 | |
| | Cairns | RMP003 | RNA | H6 | -16.919 | 145.778 | |
| | Cairns | RMP004 | RNA | H6 | -16.919 | 145.778 | |
| | Cairns | RMP005 | RNA | H6 | -16.919 | 145.778 | |
| | Cairns | RMP006 | RNA | H37 | -16.919 | 145.778 | |
| | Cairns | RMP007 | RNA | H44 | -16.919 | 145.778 | |
| | Cairns | RMP009 | RNA | H6 | -16.919 | 145.778 | |
| | Cairns | RMP010 | RNA | H6 | -16.919 | 145.778 | |
| | Cairns | RMP012 | RNA | H37 | -16.919 | 145.778 | |
| | Cairns | RMP015 | RNA | H45 | -16.919 | 145.778 | |
| | Cairns | RMP016 | RNA | H37 | -16.919 | 145.778 | |
| | Cairns | RMP020 | RNA | H46 | -16.919 | 145.778 | |
| | Croydon | RM299 | RNA | H37 | -18.206 | 142.240 | |
| | Croydon | RM300 | RNA | H6 | -18.206 | 142.240 | |
| | Croydon | RM302 | RNA | H6 | -18.206 | 142.240 | |
| | Croydon | RM304 | RNA | H38 | -18.206 | 142.240 | |
| | Daintree | RM399 | RNA | H41 | -16.250 | 145.317 | This study |
| | Daintree | RM400 | RNA | H6 | -16.250 | 145.317 | |
| | Daintree | RM401 | RNA | H29 | -16.250 | 145.317 | |
| | Daintree | RM402 | RNA | H42 | -16.250 | 145.317 | |
| | Daintree | RM403 | RNA | H37 | -16.250 | 145.317 | |
| | Daintree | RM404 | RNA | H37 | -16.250 | 145.317 | |
| | Gordonvale | RM260 | RNA | H6 | -17.083 | 145.796 | |
| | Gordonvale | RM261 | RNA | H34 | -17.083 | 145.796 | |
| | Gordonvale | RM262 | RNA | H35 | -17.083 | 145.796 | |
| | Gordonvale | RM265 | WGS | H6 | -17.083 | 145.796 | |
| | Gordonvale | RM278 | RNA | H36 | -17.083 | 145.796 | |
| | Gordonvale | RM280 | RNA | H21 | -17.083 | 145.796 | |
| | Innisfail | RM106 | RNA | H26 | -17.496 | 146.047 | |
| | Innisfail | RM108 | RNA | H27 | -17.496 | 146.047 | |
| | Innisfail | RM118 | RNA, WGS | H6 | -17.496 | 146.047 | |
| | Innisfail | RM127 | RNA, WGS | H28 | -17.496 | 146.047 | |
| | Innisfail | RM135 | RNA | H27 | -17.496 | 146.047 | |
| | Rossville | RM169 | RNA | H29 | -15.705 | 145.223 | |
| | Rossville | RM170 | RNA | H30 | -15.705 | 145.223 | |
| | Rossville | RM171 | RNA | H31 | -15.705 | 145.223 | |
| | Rossville | RM179 | RNA | H32 | -15.705 | 145.223 | |
| | Rossville | RM189 | RNA | H33 | -15.705 | 145.223 | |
| | Mossman | | | | | | |
| | Rockhampton | | | | | | Slade and Moritz et al. |
| | Brisbane | | | | | | |
| New South Wales | | | | | | | |
| | Lennox Head | | | | | | Slade and Moritz et al. |
| **East of Andes** | | KP704699 - KP704700 | | | | | |
| | | KP979778 | | | | | |
| | | KP979779 - KP979781 | | | | | |
| | | KP704701 - KP704713 | | | | | |
| | | KP704714 - KP704723 | | | | | |
| | | KP704724 | | | | | |
| | | KP704725 - KP704734 | | | | | Acevedo et al. |
| **West of Andes** | | KP704669 - KP704677 | | | | | |
| | | KP704678 - KP704684 | | | | | |
| | | KP704685 - KP704691 | | | | | |
| | | KP704692 - KP704693 | | | | | |
| | | KP704694 - KP704696 | | | | | |
| | | KP704697 - KP704698 | | | | | |

**Table S2: Mitogenome annotation of *Rhinella marina* with reference genome and the number of parsimony informative sites and singleton sites within 132 individuals.**

| Gene | Start | Stop | Strand (+/-) | Length | Parsimony informative site | Singleton variable site |
|---|---|---|---|---|---|---|
| tRNA(L1) | 1 | 72 | + | 72 | 0 | 0 |
| tRNA(T) | 73 | 144 | + | 72 | 0 | 0 |
| tRNA(P) | 144 | 212 | - | 69 | 0 | 0 |
| tRNA(F) | 212 | 279 | + | 68 | 0 | 0 |
| 16S rRNA | 280 | 1214 | + | 935 | 5 | 3 |
| tRNA(V) | 1212 | 1280 | + | 69 | 0 | 0 |
| 23S rRNA | 1281 | 2885 | + | 1605 | 13 | 6 |
| tRNA(L2) | 2885 | 2957 | + | 73 | 0 | 0 |
| NAD1 | 2973 | 3911 | + | 939 | 18 | 6 |
| tRNA(I) | 3919 | 3989 | + | 71 | 0 | 0 |
| tRNA(Q) | 3989 | 4059 | - | 71 | 0 | 0 |
| tRNA(M) | 4059 | 4127 | + | 69 | 0 | 0 |
| NAD2 | 4128 | 5156 | + | 1029 | 11 | 4 |
| tRNA(W) | 5161 | 5230 | + | 70 | 0 | 0 |
| tRNA(A) | 5231 | 5299 | - | 69 | 1 | 0 |
| tRNA(N) | 5300 | 5372 | - | 73 | 0 | 0 |
| tRNA(C) | 5401 | 5464 | - | 64 | 2 | 0 |
| tRNA(Y) | 5465 | 5534 | - | 70 | 0 | 0 |
| COX1 | 5542 | 7068 | + | 1527 | 20 | 4 |
| tRNA(S2) | 7082 | 7152 | - | 71 | 0 | 0 |
| tRNA(D) | 7154 | 7222 | + | 69 | 1 | 0 |
| COX2 | 7224 | 7895 | + | 672 | 5 | 0 |
| tRNA(K) | 7912 | 7983 | + | 72 | 1 | 0 |
| ATP8 | 7985 | 8143 | + | 159 | 2 | 2 |
| ATP6 | 8140 | 8817 | + | 678 | 10 | 4 |
| COX3 | 8823 | 9605 | + | 783 | 7 | 5 |
| tRNA(G) | 9607 | 9676 | + | 70 | 0 | 0 |
| NAD3 | 9674 | 10015 | + | 342 | 7 | 3 |
| tRNA(R) | 10017 | 10085 | + | 69 | 1 | 0 |
| NAD4L | 10125 | 10382 | + | 258 | 4 | 0 |
| NAD4-0 | 10379 | 11737 | + | 1359 | 16 | 4 |
| tRNA(H) | 11744 | 11812 | + | 69 | 0 | 0 |
| tRNA(S1) | 11813 | 11879 | + | 67 | 1 | 0 |
| NAD5 | 11933 | 13705 | + | 1773 | 20 | 7 |
| NAD6 | 13707 | 14198 | - | 492 | 5 | 0 |
| tRNA(E) | 14199 | 14266 | - | 68 | 0 | 0 |
| COB | 14271 | 15401 | + | 1131 | 20 | 4 |
| Control region | 15402 | 18152 | | 2751 | 27 | 11 |
| Intergenic | | | | | 1 | 1 |
| | | | | **Total** | **198** | **64** |

**Table S3: Summary of the complete mitogenomes of 203 Anuran and 5 non-Anuran as outgroup downloaded from NCBI BioProject database for phylogenetic tree construction.**

| Accession number | Order | Organism |
|---|---|---|
| NC_066225.1 | Anura | *Rhinella marina* |
| NC_001573.1 | Anura | *Xenopus laevis* |
| NC_002805.1 | Anura | *Rana nigromaculata* |
| NC_005055.1 | Anura | *Fejervarya limnocharis* |
| NC_005794.2 | Anura | *Bufo melanostictus* |
| NC_006402.1 | Anura | *Bombina fortinuptialis* |
| NC_006403.1 | Anura | *Hyla chinensis* |
| NC_006405.1 | Anura | *Kaloula pulchra* |
| NC_006406.1 | Anura | *Microhyla heymonsi* |
| NC_006688.1 | Anura | *Alytes obstetricans pertinax* |
| NC_006689.1 | Anura | *Bombina orientalis* |
| NC_006690.1 | Anura | *Discoglossus galganoi* |
| NC_006839.1 | Anura | *Xenopus tropicalis* |
| NC_007178.1 | Anura | *Rhacophorus schlegelii* |
| NC_007440.2 | Anura | *Limnonectes fujianensis* |
| NC_007888.1 | Anura | *Mantella madagascariensis* |
| NC_008144.1 | Anura | *Pelobates cultripes* |
| NC_008410.1 | Anura | *Bufo gargarizans* |
| NC_009258.1 | Anura | *Bombina variegata* |
| NC_009264.1 | Anura | *Rana plancyi* |
| NC_009422.1 | Anura | *Microhyla ornata* |
| NC_009423.1 | Anura | *Amolops tormotus* |
| NC_009886.1 | Anura | *Bufo japonicus* |
| NC_010232.1 | Anura | *Dryophytes japonicus* voucher IABHU6123 |
| NC_010233.1 | Anura | *Microhyla okinavensis* |
| NC_011049.1 | Anura | *Bombina maxima* |
| NC_012647.1 | Anura | *Fejervarya cancrivora* |
| NC_013270.1 | Anura | *Quasipaa spinosa* |
| NC_014581.1 | Anura | *Hoplobatrachus tigerinus* |
| NC_014584.1 | Anura | *Euphlyctis hexadactylus* |
| NC_014685.1 | Anura | *Occidozyga martensii* |
| NC_014691.1 | Anura | *Leiopelma archeyi* |
| NC_015305.1 | Anura | *Rana ishikawae* |
| NC_015615.1 | Anura | *Hymenochirus boettgeri* |
| NC_015617.1 | Anura | *Pipa carvalhoi* |
| NC_015618.1 | Anura | *Pseudhymenochirus merlini* |
| NC_015620.1 | Anura | *Rhinophrynus dorsalis* |
| NC_016119.1 | Anura | *Nanorana pleskei* |
| NC_018771.1 | Anura | *Babina adenopleura* |
| NC_018775.1 | Anura | *Xenopus victorianus* |
| NC_018776.1 | Anura | *Xenopus borealis* |
| NC_018785.1 | Anura | *Atympanophrys shapingensis* |
| NC_019615.1 | Anura | *Hoplobatrachus rugulosus* |
| NC_019998.1 | Anura | *Heleophryne regis* |
| NC_019999.1 | Anura | *Lechriodus melanopyga* |
| NC_020000.1 | Anura | *Pelodytes cf. punctatus II-2011* |
| NC_020001.1 | Anura | *Sooglossus thomasseti* |
| NC_020002.1 | Anura | *Telmatobius bolivianus* |
| NC_020044.1 | Anura | *Kaloula borealis* |
| NC_020048.1 | Anura | *Bufo tibetanus* |
| NC_020610.1 | Anura | *Leptolalax oshanensis* |
| NC_021476.1 | Anura | *Bombina microdeladigitora* voucher 1770 |
| NC_021477.1 | Anura | *Bombina lichuanensis* voucher BL3589 |
| NC_021937.1 | Anura | *Quasipaa boulengeri* |
| NC_022696.1 | Anura | *Rana catesbeiana* |
| NC_022870.1 | Anura | *Babina holsti* |
| NC_022871.1 | Anura | *Babina subaspera* |
| NC_022872.1 | Anura | *Babina okinavana* |
| NC_023379.1 | Anura | *Breviceps adspersus* isolate: No. B01 |
| NC_023380.1 | Anura | *Hemisus marmoratus* |
| NC_023381.1 | Anura | *Hyperolius marmoratus* |
| NC_023382.1 | Anura | *Trichobatrachus robustus* isolate: No. B05 |
| NC_023528.1 | Anura | *Rana dybowskii* |
| NC_023529.1 | Anura | *Rana cf. chensinensis* QYCRCCH_C |
| NC_023949.1 | Anura | *Amolops ricketti* |
| NC_024180.1 | Anura | *Amolops mantzorum* |
| NC_024272.1 | Anura | *Nanorana taihangnica* |
| NC_024427.1 | Anura | *Leptobrachium boringii* voucher SCUM120630 |
| NC_024547.1 | Anura | *Microhyla pulchra* |
| NC_024548.1 | Anura | *Rana kunyuensis* |
| NC_024603.1 | Anura | *Odorrana margaretae* |
| NC_024748.1 | Anura | *Hylarana guentheri* |
| NC_024843.1 | Anura | *Quasipaa yei* voucher HNNU0908I061 |
| NC_025226.1 | Anura | *Glandirana tientaiensis* |
| NC_025309.1 | Anura | *Hyla annectans* |
| NC_025575.1 | Anura | *Pelophylax cretensis* isolate CR03 |

| | | |
|---|---|---|
| NC_025591.1 | Anura | *Amolops wuyiensis* |
| NC_026524.1 | Anura | *Hyla tsinlingensis* voucher HTSIN20141129 |
| NC_026789.1 | Anura | *Nanorana parkeri* |
| NC_026893.1 | Anura | *Pelophylax cypriensis* isolate GM157-11 |
| NC_026894.1 | Anura | *Pelophylax epeiroticus* isolate MPFC1392 |
| NC_026895.1 | Anura | *Pelophylax kurtmuelleri* isolate MPFC1475 |
| NC_026896.1 | Anura | *Pelophylax shqipericus* isolate GM1013 |
| NC_027072.1 | Anura | *Leiopelma hochstetteri* |
| NC_027236.1 | Anura | *Rana sylvatica* |
| NC_027452.1 | Anura | *Rhacophorus dennysi* |
| NC_027671.1 | Anura | *Platymantis vitianus* |
| NC_027686.1 | Anura | *Bufo stejnegeri* voucher y-d20130012 |
| NC_027827.1 | Anura | *Odorrana schmackeri* |
| NC_028283.1 | Anura | *Rana okaloosae* voucher LodgeLab Rokaloosae_1 |
| NC_028296.1 | Anura | *Rana draytonii* voucher LodgeLab Rdraytonii_1 |
| NC_028424.1 | Anura | *Bufotes raddei* |
| NC_028521.1 | Anura | *Rana huanrensis* voucher y-d20130058 |
| NC_029196.1 | Anura | *Pelophylax cf. bedriagae* AFL1 |
| NC_029197.1 | Anura | *Pelophylax cf. bedriagae* MPFC1082 |
| NC_029198.1 | Anura | *Pelophylax cf. terentievi GM87-239* |
| NC_029199.1 | Anura | *Pelophylax cf. terentievi* MPFC1736 |
| NC_029200.1 | Anura | *Pelophylax bedriagae* isolate GM-J0142 |
| NC_029201.1 | Anura | *Pelophylax cf. bedriagae* GM178 |
| NC_029250.1 | Anura | *Amolops loloensis* voucher SM-ZDTW-01 |
| NC_029409.1 | Anura | *Kaloula rugifera* |
| NC_029410.1 | Anura | *Hyla ussuriensis* voucher HRB1506014 |
| NC_029754.1 | Anura | *Fejervarya multistriata* voucher zjbj3 |
| NC_030042.1 | Anura | *Rana amurensis* voucher SYNU11100268 |
| NC_030049.1 | Anura | *Microhyla butleri* isolate CIbSZ20150408 |
| NC_030054.1 | Anura | *Anomaloglossus baeobatrachus* voucher AF2590 |
| NC_030211.1 | Anura | *Rugosa emeljanovi* voucher HRB1506079 |
| NC_030333.1 | Anura | *Telmatobius chusmisensis* |
| NC_030605.1 | Anura | *Oreolalax major* |
| NC_030627.1 | Anura | *Oreolalax lichuanensis* |
| NC_031411.1 | Anura | *Leptobrachium leishanense* |
| NC_031426.1 | Anura | *Scutiger ningshanensis* |
| NC_032380.1 | Anura | *Dryophytes suweonensis* |
| NC_034983.1 | Anura | *Odorrana wuchuanensis* |
| NC_034984.1 | Anura | *Odorrana hainanensis* |
| NC_035803.1 | Anura | *Rana chaochiaoensis* voucher SM-ZJLW-01 |
| NC_035804.1 | Anura | *Rana kukunoris* voucher RG-GYLW-01 |
| NC_035805.1 | Anura | *Rana omeimontis* voucher YC-EMLW-01 |
| NC_036493.1 | Anura | *Bokermannohyla alvarengai* |
| NC_037377.1 | Anura | *Scaphiopus holbrookii* |
| NC_037378.1 | Anura | *Melanophryniscus moreirae* |
| NC_037379.1 | Anura | *Hyloxalus subpunctatus* |
| NC_037380.1 | Anura | *Phyllobates terribilis* |
| NC_037382.1 | Anura | *Oreolalax multipunctatus* voucher CIB2013wb091 |
| NC_037854.1 | Anura | *Anomaloglossus surinamensis* voucher AF0585 |
| NC_037855.1 | Anura | *Anomaloglossus dewynteri* voucher PG660 |
| NC_037856.1 | Anura | *Anomaloglossus degranvillei* voucher PG601 |
| NC_037857.1 | Anura | *Anomaloglossus blanci* voucher AF0932 |
| NC_038130.1 | Anura | *Microhyla mixtura* |
| NC_039094.1 | Anura | *Nanorana ventripunctata* |
| NC_039176.1 | Anura | *Microhyla taraiensis* |
| NC_039411.1 | Anura | *Kaloula verrucosa* |
| NC_039758.1 | Anura | *Mantella baroni* |
| NC_041426.1 | Anura | *Pseudis tocantins* |
| NC_042226.1 | Anura | *Rana temporaria* |
| NC_042258.1 | Anura | *Hoplobatrachus chinensis x Hoplobatrachus rugulosus* |
| NC_042501.1 | Anura | *Bombina bombina* |
| NC_042797.1 | Anura | *Polypedates braueri* |
| NC_043768.1 | Anura | *Odorrana livida* |
| NC_043955.1 | Anura | *Polypedates megacephalus* voucher 20130003 |
| NC_044480.1 | Anura | *Pyxicephalus adspersus* |
| NC_044865.1 | Anura | *Xenopus calcaratus* isolate R7934_NMP6V_74630.1 |
| NC_044866.1 | Anura | *Xenopus mellotropicalis* isolate R7933_AMNH17288 |
| NC_044867.1 | Anura | *Xenopus epitropicalis* isolate R7932_AMNH17278 |
| NC_044868.1 | Anura | *Xenopus largeni* isolate R7939_MHNG2644.059 |
| NC_044869.1 | Anura | *Xenopus petersii* isolate R7944_MHNG2645.094 |
| NC_044870.1 | Anura | *Xenopus poweri* isolate R7946_MHNG2645.073 |
| NC_044871.1 | Anura | *Xenopus gilli* isolate R7943_Xr_0_3 |
| NC_044872.1 | Anura | *Xenopus parafraseri* isolate R7942_MHNG2645.084 |
| NC_044873.1 | Anura | *Xenopus pygmaeus* isolate R7940_AMNH17320 |
| NC_044874.1 | Anura | *Xenopus allofraseri* isolate R7941_MCZHerpetology_A_148164 |
| NC_044875.1 | Anura | *Xenopus wittei* isolate R7949_MHNG2645.092 |
| NC_044876.1 | Anura | *Xenopus amieti* isolate R7948_MHNG2645.079 |
| NC_044877.1 | Anura | *Xenopus boumbaensis* isolate R7950_MHNG2644.057 |
| NC_044878.1 | Anura | *Xenopus andrei* isolate R7951_MHNG2645.072 |

| | | |
|---|---|---|
| NC_044879.1 | Anura | *Xenopus itombwensis* isolate R7952_MCZHerpetology_A_138192 |
| NC_044880.1 | Anura | *Xenopus lenduensis* isolate R7953_MCZHerpetology_A_140111 |
| NC_044881.1 | Anura | *Xenopus vestitus* isolate R7954_RT2 |
| NC_044882.1 | Anura | *Xenopus ruwenzoriensis* isolate R7956_AMNH17317 |
| NC_044883.1 | Anura | *Xenopus kobeli* isolate R7957_MCZHerpetology_A_148037 |
| NC_044884.1 | Anura | *Xenopus eysoole* isolate R7958_MCZHerpetology_A_148095 |
| NC_044885.1 | Anura | *Xenopus longipes* isolate R7959_MCZHerpetology_A_148072 |
| NC_044886.1 | Anura | *Xenopus clivii* isolate R7935_MHNG2645.069 |
| NC_044887.1 | Anura | *Xenopus muelleri* isolate R7936_Xen203 |
| NC_044888.1 | Anura | *Xenopus fischbergi* isolate R7938_AMNH17297 |
| NC_044901.1 | Anura | *Amolops granulosus* |
| NC_045110.1 | Anura | *Microhyla fissipes* |
| NC_045917.1 | Anura | *Dryophytes versicolor* |
| NC_046047.1 | Anura | *Bufotes zamdaensis* |
| NC_046387.1 | Anura | *Rhacophorus omeimontis* |
| NC_047224.1 | Anura | *Anaxyrus americanus* |
| NC_047225.1 | Anura | *Bufotes pewzowi* |
| NC_049862.1 | Anura | *Oreolalax omeimontis* |
| NC_050665.1 | Anura | *Bufotes variabilis* isolate DM1061 |
| NC_050884.1 | Anura | *Odorrana graminea* |
| NC_051953.1 | Anura | *Pelobates fuscus* isolate DM1190 |
| NC_053712.1 | Anura | *Odorrana exiliversabilis* voucher LSU20200716WY02 |
| NC_056269.1 | Anura | *Quasipaa exilispinosa* voucher LSU20200418001WY |
| NC_056272.1 | Anura | *Rana uenoi* |
| NC_056343.1 | Anura | *Oreolalax schmidti* |
| NC_056366.1 | Anura | *Leptodactylus fallax* |
| NC_057198.1 | Anura | *Hylarana latouchii* voucher LSU20200422001ZL |
| NC_057468.1 | Anura | *Leptobrachium liui* voucher LSU20191108JLS001 |
| NC_057992.1 | Anura | *Phrynoglossus myanhessei* voucher SMF 103841 |
| NC_058599.1 | Anura | *Rana johnsi* |
| NC_059861.1 | Anura | *Odorrana grahami* |
| NC_060306.1 | Anura | *Rana dabieshanensis* |
| NC_060625.1 | Anura | *Megophrys sp.* HW-2022 |
| NC_061370.1 | Anura | *Rana longicrus* voucher LSU20210423MXZY001 |
| NC_061371.1 | Anura | *Rana hanluica* voucher LSU20201009LSLD001 |
| NC_061389.1 | Anura | *Pipa pipa* |
| NC_061400.1 | Anura | *Gracixalus yunnanensis* |
| NC_061925.1 | Anura | *Pipa snethlageae* |
| NC_061926.1 | Anura | *Pipa myersi* |
| NC_062077.1 | Anura | *Bufotes turanensis* |
| NC_062326.1 | Anura | *Hyla sanchiangensis* voucher LSU20210421MXHT001 |
| NC_062354.1 | Anura | *Polypedates impresus* |
| NC_062355.1 | Anura | *Polypedates mutus* |
| NC_062356.1 | Anura | *Polypedates leucomystax* |
| NC_062878.1 | Anura | *Rhacophorus chenfui* |
| NC_063648.1 | Anura | *Dryophytes andersonii* isolate ARW103 |
| NC_063649.1 | Anura | *Dryophytes femoralis* isolate ARW436 |
| NC_065297.1 | Anura | *Odorrana jingdongensis* |
| AY954504.1 | Caeciliidae | *Boulengerula taitanus* |
| NC_006303.1 | Caeciliidae | *Rhinatrema bivittatum* |
| JX508764.1 | Caudata | *Salamandrella keyserlingii* isolate SKN9 |
| KX298241.1 | Caudata | *Andrias davidianus* isolate HZ voucher HS16093 |
| NC_006889.1 | Caudata | *Ambystoma dumerilii* |

**Table S4: Comparison between WGS and RNASeq-derived mitogenomes.**

| Sample | Position | Region | WGS | RNASeq | Conversion |
|---|---|---|---|---|---|
| RMF048 | 267 | tRNA(F) | A | G | Transition |
| | 2244 | 23S rRNA | A | T | Transversion |
| | 15524 | control region | C | T | Transversion |
| | 15705 | | T | C | Transition |
| | 15752 | | C | T | Transition |
| | 15929 | | C | T | Transition |
| | 15933 | | T | C | Transition |
| | 15975 | | T | C | Transition |
| | 16011 | | A | G | Transition |
| | 16115 | | A | G | Transition |
| | 16211 | | A | G | Transition |
| | 16775 | | G | A | Transition |
| | 16916 | | T | C | Transition |
| | 17226 | | C | T | Transition |
| RMH006 | 2244 | 23S rRNA | A | T | Transversion |
| RMH051 | 2244 | 23S rRNA | A | T | Transversion |
| RMH052 | 2244 | 23S rRNA | A | T | Transversion |
| RM0010 | 477 | 16S rRNA | C | T | Transition |
| | 2244 | 23S rRNA | A | T | Transversion |
| | 5162 | tRNA(W) | G | A | Transition |
| | 7162 | tRNA(D) | A | T | Transversion |
| | 7920 | tRNA(K) | A | T | Transversion |
| RM0118 | 2244 | 23S rRNA | A | T | Transversion |
| | 7162 | tRNA(D) | A | G | Transition |
| | 7920 | tRNA(K) | A | G | Transition |
| RM0127 | 2244 | 23S rRNA | A | T | Transversion |
| | 5162 | tRNA(W) | G | A | Transition |
| | 7162 | tRNA(D) | A | T | Transversion |
| | 7920 | tRNA(K) | A | G | Transition |
| RM0045 | 2244 | 23S rRNA | A | T | Transversion |
| | 5162 | tRNA(W) | G | A | Transition |
| | 7162 | tRNA(D) | A | G | Transition |
| | 7920 | tRNA(K) | A | G | Transition |

**Table S5: NUMTs blocks in the cane toad nuclear genome with respect to its mtDNA.**

| Contig Name | Contig Start | Contig End | Strand | mtDNA Fragment | Fragment Length | Matches | E Value |
|---|---|---|---|---|---|---|---|
| ctg10165_RHIMB__RM170330.10165 | 170828 | 170875 | - | 17943-17990 | 48 | 42 | 5.00E-05 |
| ctg104_RHIMB__RM170330.104 | 134299 | 134336 | - | 17943-17980 | 38 | 36 | 5.00E-05 |
| ctg11815_RHIMB__RM170330.11815 | 194991 | 195051 | - | 17929-17986 | 61 | 52 | 1.00E-05 |
| ctg1194_RHIMB__RM170330.1194 | 18743 | 18780 | + | 17943-17980 | 38 | 36 | 5.00E-05 |
| ctg1208_RHIMB__RM170330.1208 | 498924 | 498989 | - | 17928-17990 | 66 | 55 | 3.00E-07 |
| ctg1221_RHIMB__RM170330.1221 | 675608 | 675653 | - | 17923-17968 | 46 | 41 | 5.00E-05 |
| ctg12503_RHIMB__RM170330.12503 | 78151 | 78290 | - | 17944-18084 | 140 | 107 | 1.00E-05 |
| ctg12723_RHIMB__RM170330.12723 | 19348 | 19385 | + | 17931-17968 | 38 | 36 | 5.00E-05 |
| ctg1367_RHIMB__RM170330.1367 | 430982 | 431026 | + | 17933-17978 | 45 | 42 | 1.00E-05 |
| ctg13991_RHIMB__RM170330.13991 | 218571 | 218608 | + | 17943-17980 | 38 | 36 | 5.00E-05 |
| ctg15571_RHIMB__RM170330.15571 | 25520 | 25559 | - | 17943-17981 | 40 | 38 | 5.00E-05 |
| ctg16362_RHIMB__RM170330.16362 | 46563 | 46600 | + | 17931-17968 | 38 | 36 | 5.00E-05 |
| ctg16795_RHIMB__RM170330.16795 | 20291 | 20334 | + | 17936-17980 | 44 | 41 | 5.00E-05 |
| ctg18469_RHIMB__RM170330.18469 | 54795 | 54837 | - | 17932-17974 | 43 | 39 | 5.00E-05 |
| ctg19869_RHIMB__RM170330.19869 | 28837 | 28882 | + | 17943-17988 | 46 | 41 | 5.00E-05 |
| ctg23132_RHIMB__RM170330.23132 | 89407 | 89455 | - | 17943-17988 | 49 | 45 | 5.00E-05 |
| ctg24175_RHIMB__RM170330.24175 | 26491 | 26532 | + | 17939-17979 | 42 | 40 | 4.00E-06 |
| ctg24322_RHIMB__RM170330.24322 | 38880 | 38929 | + | 17943-17990 | 50 | 45 | 5.00E-05 |
| ctg2588_RHIMB__RM170330.2588 | 91193 | 91240 | + | 17943-17990 | 48 | 42 | 5.00E-05 |
| ctg2649_RHIMB__RM170330.2649 | 238602 | 238662 | + | 18024-18087 | 61 | 53 | 5.00E-05 |
| ctg26654_RHIMB__RM170330.26654 | 578 | 622 | - | 17933-17980 | 45 | 43 | 1.00E-05 |
| ctg26_RHIMB__RM170330.26 | 291650 | 291714 | + | 17944-18005 | 65 | 53 | 1.00E-05 |
| ctg29742_RHIMB__RM170330.29742 | 34189 | 34226 | + | 17943-17980 | 38 | 36 | 5.00E-05 |
| ctg30050_RHIMB__RM170330.30050 | 4952 | 4995 | - | 17939-17979 | 44 | 40 | 5.00E-05 |
| ctg31312_RHIMB__RM170330.31312 | 16554 | 16604 | - | 17941-17990 | 51 | 45 | 1.00E-05 |
| ctg3490_RHIMB__RM170330.3490 | 114481 | 114529 | - | 17944-17989 | 49 | 43 | 5.00E-05 |
| ctg35_RHIMB__RM170330.35 | 2052238 | 2052282 | + | 17944-17987 | 45 | 41 | 5.00E-05 |
| ctg3738_RHIMB__RM170330.3738 | 186479 | 186519 | + | 17943-17983 | 41 | 38 | 5.00E-05 |
| ctg3992_RHIMB__RM170330.3992 | 102790 | 102860 | + | 17929-17994 | 71 | 58 | 5.00E-05 |
| ctg427_RHIMB__RM170330.427 | 45564 | 45608 | - | 17943-17985 | 45 | 41 | 5.00E-05 |
| ctg4517_RHIMB__RM170330.4517 | 64852 | 64896 | - | 17930-17976 | 45 | 42 | 5.00E-05 |
| ctg4787_RHIMB__RM170330.4787 | 26493 | 26542 | + | 17931-17979 | 50 | 44 | 5.00E-05 |
| ctg5197_RHIMB__RM170330.5197 | 126698 | 126745 | + | 17943-17989 | 48 | 43 | 5.00E-05 |
| ctg5682_RHIMB__RM170330.5682 | 124532 | 124576 | - | 17936-17979 | 45 | 41 | 5.00E-05 |
| ctg626_RHIMB__RM170330.626 | 23009 | 23057 | - | 17943-17990 | 49 | 44 | 1.00E-05 |
| ctg6533_RHIMB__RM170330.6533 | 64253 | 64301 | + | 17942-17991 | 49 | 44 | 5.00E-05 |
| ctg6731_RHIMB__RM170330.6731 | 62271 | 62318 | + | 17943-17990 | 48 | 42 | 5.00E-05 |
| ctg7238_RHIMB__RM170330.7238 | 69867 | 69912 | + | 17943-17987 | 46 | 42 | 1.00E-05 |
| ctg8177_RHIMB__RM170330.8177 | 10312 | 10370 | + | 17931-17988 | 59 | 50 | 1.00E-05 |
| ctg8271_RHIMB__RM170330.8271 | 35978 | 36037 | - | 18028-18087 | 60 | 50 | 4.00E-06 |
| ctg8379_RHIMB__RM170330.8379 | 8554 | 8601 | + | 17943-17988 | 48 | 43 | 1.00E-05 |
| ctg8953_RHIMB__RM170330.8953 | 85556 | 85601 | - | 17943-17988 | 46 | 41 | 5.00E-05 |
